# Synthetic bacterial community colonizes wheat roots grown in soil and mimics the assembly pattern of a field community

**DOI:** 10.1101/2025.10.01.677727

**Authors:** Frederik Bak, Jakob Klinge Meier, Bo Markussen, Kitzia Yashvelt Molina-Zamudio, Clara Tang, Mette H. Nicolaisen

## Abstract

The root microbiome is important for plant development. The impact of the root microbiome is the sum of multiple complex interactions among microorganisms, the plant and the environment. This complexity can be reduced by designing synthetic bacterial communities (SynComs) consisting of bacteria isolated from the roots, making it possible to study these interactions. However, the translational power from SynCom experiments to explain field observations is still very low, which demonstrates the need for development of SynComs that colonize plants comparable to what is observed in the field. Hence, we developed a SynCom consisting of 13 different strains from 13 genera with varying phenotypes originating from the roots of winter wheat (*Triticum aestivum* cv. Sheriff). The SynCom was inoculated into gamma-irradiated soil prior to sowing and community assembly was determined over 4 weeks using 16S rRNA amplicon sequencing. The SynCom supported growth of winter wheat over a 4-week period and developed in a comparable manner to a more diverse natural community (NatCom) obtained from a soil solution. Furthermore, the temporal dynamics of the majority of the SynCom strains mimicked the development in relative abundance of their respective genera in field grown winter wheat of similar cultivar. However, this could not be translated to a different cultivar (Herup). Our results demonstrate how SynComs inoculated into gamma-irradiated soil can provide a framework for bridging the gap between greenhouse and lab experiments and field observations. At the same time it highlights the plant-genotype specific impact on community assembly.

## Introduction

Microorganisms interact with plants above- and belowground and are important for plant health and development, as they provide beneficial functions such as nutrient acquisition, disease suppression, or reduction of abiotic and biotic stresses (Mendes *et al*., 2013; Vandenkoornhuyse *et al*., 2015; Compant *et al*., 2019). However, the processes governing root microbial community assembly are not fully understood, despite an enormous effort being made to advance this field (Trivedi *et al*., 2020). In the rhizosphere, this is primarily due to the high chemical complexity and microbial diversity.

The seed carries the starting members of the rhizosphere microbiome, yet the largest proportion originates from the soil. The initial assembly of the rhizosphere community assembly is primarily driven by stochastic effects such as dispersal and drift (Dini-Andreote and Raaijmakers, 2018). While the variability in soil microbiomes lead to differences in the microbiome of the same plant species across soil types, deterministic processes are becoming more important for community assembly at later developmental stages (Dini-Andreote and Raaijmakers, 2018; Trivedi *et al*., 2020; Rüger *et al*., 2021). However, our knowledge on microbial community development is primarily based on observational studies that offer limited information on the causal effects. To leverage our understanding of root microbiome assembly and establish causal relationships, it is crucial to develop and use reproducible model systems with reduced complexity (Vorholt *et al*., 2017; Northen *et al*., 2024), which still reflect the actual soil systems. Such model systems include synthetic communities (SynComs) which provide the possibility to test hypotheses on causality in the rhizosphere (Vorholt *et al*., 2017).

Until now, studies applying SynComs for understanding root-microbe interactions are performed solely in media (Ma *et al*., 2021), gnotobiotic systems (Harbort *et al*., 2020; Wippel *et al*., 2021; Lin *et al*., 2024) or artificial soil mixtures (Zhou *et al*., 2022), whereas real soil systems have not been used. These SynCom studies have been used to elucidate plant-microbe interactions on *Arabidopsis thaliana* (Carlström *et al*., 2019; Wippel *et al*., 2021; Pfeilmeier *et al*., 2024), *Lotus japonicus* (Wippel *et al*., 2021), maize (Niu *et al*., 2017) and tomato (Zhou *et al*., 2022). These have revealed interesting results of priority effects (Carlström *et al*., 2019; Wippel *et al*., 2021), the importance of bacterial-fungal interaction in pathogen suppression (Zhou *et al*., 2022), and the identification of *Erwinia cloacea* as a keystone species in the maize rhizosphere (Niu *et al*., 2017). Still, the complexity of soil with varying niches, physical hindering for motility and chemical gradients are omitted, questioning the translational power of the findings to the field. Hence, the use of soil in SynCom experiments can help bridge the gap in translational research (Sessitsch *et al*., 2019).

There are multiple strategies for selecting the members and size of a SynCom (Jing *et al*., 2024) and they can vary in size from a few members, e.g. 7 strains (Niu *et al*., 2017), up to more than 100 strains (Pfeilmeier *et al*., 2024), and from one or more kingdoms (Zhou *et al*., 2022). While the SynComs do not represent the full diversity of a given environment, individual and rhizo-compatible strains could represent a genus in studies of rhizosphere community assembly, as growth rate is an important factor governing bacterial success in rhizosphere colonization (López *et al*., 2023), and is strongly conserved at the genus level (Weissman *et al*., 2021), even though high microdiversity is found within species colonizing the rhizosphere (Herms *et al*., 2024). This is supported by the notion that core bacteria are often being classified at the genus level (Simonin *et al*., 2020), and it suggests that strains within a genus should be interchangeable in community assembly. Still, it remains to be determined if community dynamics in a SynCom can reflect those under field conditions.

To test this, we designed a SynCom that could colonize and establish a community on the plant roots and support plant growth. We hypothesized that the SynCom would follow the overall rules of root assembly with high differences in beta diversity at early stages of community assembly due to initial random assembly followed by a reduced variation in beta diversity reflecting the increasing deterministic processes. Secondly, as growth rate is conserved at the genus level, we hypothesized that the relative abundances of singles strains would be similar to the relative abundance of genera in a natural community (NatCom) or in the field under natural conditions.

We present the establishment of a root-compatible SynCom consisting of 13 bacterial strains representing 13 bacterial genera isolated from the roots of winter wheat (*Triticum aestivum cv*. Sheriff). These genera comprised both rare and core members of the wheat rhizosphere as presented previously (Simonin *et al*., 2020). We inoculated the SynCom to gamma-irradiated soil prior to wheat seedling transplantation to ensure active colonization of the bacteria to the growing roots, and compared microbial assembly of the SynCom on wheat roots with the assembly based on a NatCom and field grown wheat plants. We show that most SynCom members have similar temporal dynamics to their corresponding genus from NatCom and field communities, although this was wheat cultivar dependent. The approach applied in this study demonstrates the robustness of SynComs for studying rhizosphere community assembly and plant-microbe interactions, and helps bridge the translational gap between greenhouse and field for improved agricultural solutions.

## Materials and methods

### Strain isolation and selection

Winter wheat, *cv*. Sheriff, was grown in a growth chamber under controlled conditions as previously described by Guan *et al*. (2024). Three plants were harvested on day 142 at the flag leaf emergence stage (Feekes 8.0), and rhizoplane bacteria were isolated to generate a biobank of 1,344 bacterial isolates (Supplementary methods). Using 16S rRNA gene amplicon data from the winter wheat rhizosphere and rhizoplane (Zervas *et al*., 2022; Guan *et al*., 2024), we selected 13 bacterial strains for our synthetic community (Table S1). The SynCom strains’ abilities of phosphate solubilization, phytate degradation, nitrate reduction and siderophore production was determined (Supplementary methods).

### Soil description

We collected soil from the Long-Term Nutrient Depletion Trial (LTNDT) field in Taastrup, Denmark (55° 40′N, 12° 17′E). The field is classified as a sandy loam soil, and composed of 164 g kg^−1^ clay, 173 g kg^−1^ silt, 333 g kg^−1^ fine sand, 312 g kg^−1^ coarse sand and 17 g kg^−1^ organic matter (van der Bom *et al*., 2017). Soil was collected for the SynCom plant experiment in the spring of 2018 from the nitrogen fertilizer only (N1) treatment. After collection the soil was lightly dried exposed to gamma radiation (2 x 18 kGy, 10 MeV electron beam) at Sterigenics A/S, Denmark to avoid the release of phytotoxins through heating, and stored at room temperature in sealed plastic bags until use. This reduced the DNA concentration with 79% in the gamma-irradiated soil compared to untreated field soil (Table S2), and resulted in a bacterial community primarily consisting of *Streptomyces* and *Bacillus* (Fig. S1). A population of 3.0 * 10^5^ CFU/g soil was detected post gamma-irradiation.

### SynCom plant experiment

Wheat plants were grown in PVC pots as described in Guan *et al*. (2024) (Fig. S2). Gamma irradiated soil (sieved 4 mm) was mixed with sand of grain size of 0.4 - 0.9 mm (Dansand, Denmark) in a 3:2 soil:sand ratio. The sand was dried for 24 hours at 105°C and autoclaved twice at 121°C for 15 minutes prior to mixing. Soil samples (n =3) were collected to determine the bacterial communities in the starting soil, and stored at -20°C until DNA extraction. Soil moisture content was adjusted to 17% w/v with autoclaved MilliQ water. We added 0.79 ml liquid 3-1-4 NPK fertilizer (Park® Drivhusgødning, Germany) to each pot by diluting it in the water used to adjust moisture content.

Seeds of the winter wheat cultivar Sheriff were sterilized using 70% ethanol and 5% sodium hypochlorite (Acros Organics, Belgium) for three minutes each wash, followed by three rinses with sterile MilliQ water. Afterwards, we soaked the seeds in sterile MilliQ water for one hour and left them left to germinate on moist sterile filter paper at room temperature for four days in the dark prior to planting.

To prepare the SynCom inoculum, we washed overnight cultures of the SynCom strains twice, and adjusted OD_600_ to 0.6 in 0.9% sterile NaCl. We then mixed the washed cells of the 13 SynCom strains to form the final SynCom inoculum. The NatCom inoculum was created from a native soil microbial community extracted from a non-agricultural soil. We collected soil 3-5 cm below the surface from the Frederiksberg Campus (55°41’00.5”N 12°32’35.5”E, Frederiksberg, Denmark) and stored it at 5°C until use. The microbial community was extracted as in Yan *et al*. (2015) with a few modifications. Soil was sieved (4 mm) and 20 g was suspended in 190 ml 0.9% NaCl and blended 2 x 2 min. A 1,000-fold dilution was made in 0.9% NaCl to prepare the NatCom inoculum.

We added either 2.5 ml NatCom or SynCom inoculant to a two-centimeter hole in the soil in each PVC pot, and transplanted one pre-germinated seed to each pot and buried it, before adding one milliliter of sterile MilliQ. The plants were grown for 1-4 weeks in a non-sterile climate chamber with a 16/8-hour day/night cycle at 400 µmol m-^2^ s^-1^ (BX-series LED bars, Valoya, Finland). Temperatures were 20/18.5°C (day/night), with a constant 70% relative humidity. We watered the plants with sterile MilliQ every second day.

We harvested five plants from each treatment for sampling of the rhizoplane communities 7, 14, 21, and 28 days after sowing. We measured the root and shoot length from the base of the roots. We collected the rhizoplane samples as described in Guan *et al*. (2024) (Supplementary methods).

### Field experimental setup

Seeds of winter wheat cultivars Sheriff and Heerup were sown at the LTNDT field in four spatially separated plots (3x 12 m) (Fertility level: 120 kg N, 40 kg P, 240 kg K ha^−1^ y^−1^ (N_1_P_2_K_2_) fertilizer) on September 26^th^, 2023. In 2023, the field plots had only received 120 kg N ha^-1^. Each of the four plots were divided into two separate parcels, each parcel sown with several rows of one cultivar.

Nine days after sowing we sampled five plants of both cultivars from each of the four plots at random. We separated the roots and shoots at their base with a sterilized scalpel, and subjected the roots to a two-step washing process to obtain rhizoplane samples. We repeated this sampling procedure at 16, 21 and 28 Days after sowing (DAS), except only three plants per treatment were pooled for each sample instead of five, due to the larger plants. In total, we collected three bulk soil samples prior to sowing and 32 rhizoplane samples (4 time points, 2 cultivars and 4 replicates). All equipment was cleaned with 70% ethanol between each sample. The samples were stored and freeze-dried as described above.

### Bacterial community structure

We extracted DNA from all samples using the Fast-DNA™ Spin Kit for Soil (MP Biomedicals, Irvine, CA,USA) by following the manufacturer’s instructions, with minor modifications (Supplementary methods). We prepared sequencing libraries using the 799F and 1193R primers targeting the V5-V7 regions to reduce amplification of plant host DNA (Beckers *et al*., 2016). ZymoBIOMICS Microbial Community DNA Standards (Zymo Research, Irvine, CA, USA) was used to check for contamination during sequencing. Finally, indexing and sequencing was done by Eurofins Genomics (Cologne, Germany) on the Illumina MiSeq platform (2 x 300 bp).

### Amplicon processing and data analyses

Data analysis was performed using R 4.2.1 (R Core Team, 2020). Ampvis2 v.2.8.9 (Andersen *et al*., 2018) and ggplot2 v.3.5.0 packages (Wickham, 2016) were used for plotting. All of the 16S rRNA gene amplicons were processed using dada2 v.1.32 (Callahan *et al*., 2016) as detailed in (Guan *et al*., 2024). Taxonomy was assigned using the SILVA database v.138.1 (Quast *et al*., 2013). We curated the taxonomic assignment through alignment of the sequences of amplicon sequence variants (ASVs) found in the SynCom inoculum with the 16S rRNA gene sequences of the strains. In the SILVA v.138.1 database, *Peribacillus simplex* is classified as *Bacillus simplex*, and thus we corrected this in our dataset. Non-bacterial, Mitochondria and Chloroplast ASVs were removed prior to downstream analyses. Mock communities and negative controls were inspected for potential contamination.

Samples were rarefied to even depth (minimum read number =20,040 reads). This was repeated 100 times, as recommended by Schloss (2024). Subsequently, the Alpha diversity (Shannon index) was determined using the plot_richness function phyloseq v1.40 (McMurdie and Holmes, 2013). Beta diversity was calculated using Bray Curtis dissimilarities with the avgdist() function in vegan v. 2.6.4 (Oksanen *et al*., 2020), rarefying each sample to the minimum read number (20,040 reads) of rhizoplane samples 100 times (i.e., excluding bulk soil samples and controls). Principal coordinate analysis (PCoA) was performed using pco() in ecodist v.2.1.3 (Goslee and Urban, 2007). PERMANOVA was used to determine the importance of sampling time (as days after sowing) and cultivar on community composition with adonis2() in vegan, with 999 permutations and controlling for field plot using the permute v.0.9-7 (Simpson, 2022).

To determine the origin of the ASVs (i.e. inoculum or soil), we identified all ASVs from the starting inoculum or gamma-irradiated soil.. To track the SynCom strains, we subsetted the ASVs found in the SynCom inoculum from the SynCom rhizoplane samples and agglomerated them at the genus level. Upon comparison with NatCom and field samples, we extracted all ASVs belonging to the 13 SynCom genera.

### Testing for equal signs/stats on temporal dynamics

The relative abundance was analyzed separately for each genus by ANCOVA models with categorical variable Treatment (SynCom, NatCom, Field Sheriff, Field Heerup) and numerical variable (Days) as the covariate. The first time point (Days 7 or 9) was excluded due to the high dissimilarity among samples. The analyses were done on a logarithmic scale as this gave statistically valid models as validated by residual plots and normal quantile plots of the residuals. In summary, we used the statistical model

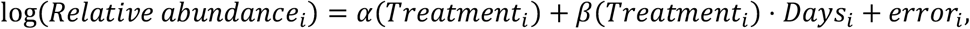

where the intercepts α(SynCom), α(NatCom), α(Field Sheriff), α(Field Heerup), the slopes β(SynCom), β(NatCom), β(Field Sheriff), β(Field Heerup), and the error variances are fitted separately for each genus. Some genera had a relative abundance of zero. As the logarithmic transformation cannot be used on zero, we estimated the ANCOVAs as Tobit models (Tobin, 1958), where a zero is interpreted as the observation that this abundance is lower than all the non-zero abundances within that genus.

To establish statistical evidence that a genus is either increasing for both SynCom and NatCom or decreasing for both SynCom and NatCom, we tested the null hypothesis that the corresponding slope parameters β(SynCom) and β(NatCom) had different signs, i.e.

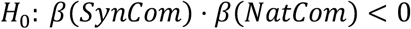

We have not been able to find a statistical test for this hypothesis. Hence, we have constructed a new test named the *Conservative One-Sided Test (COST)* to test this hypothesis. Proving validity of the COST can be done along similar lines as done for the *Two One-Sided T-test (TOST)* (Schuirmann, 1987) known from equivalence testing, but we skip the mathematical details.

COST procedure: Let 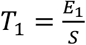 and 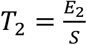 be test statistics for two scalar parameters *β*_l_ and *β*_2_ so that *E*_l_ and *E*_2_ are stochastically independent, *S* > 0 is shared, more negative *T*_*i*_ provides more evidence against *β*_*i*_ > 0, more positive *T*_*i*_ provides more evidence against *β*_*i*_ < 0, and so that the distribution of *T*_*i*_ is symmetric when the true *β*_*i*_ equals 0. Given observed test statistics *t*_l_ and *t*_2_ a p-value for the composite null hypothesis *H*_O_: *β*_l_ · *β*_2_ < 0 may be defined by 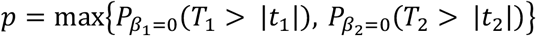 when *t*_l_ and *t*_2_ have equal signs, and *p* = 1 otherwise.

Thus, the name *Conservative One-Sided Test* arises since the COST procedure uses the larger of the two one-sided p-values when the observed signs are equal. We also note that the COST procedure can be readily changed into a test for the null hypothesis of equal signs by changing the sign of one of the samples. Moreover, the conditions for the COST procedure are fulfilled for the T-test statistics on separate slope parameters in an ANCOVA, and hence the COST p-value can easily be computed in statistical software like R by combining the p-values for the corresponding one-sided tests.

## Results

### Designing the SynCom

We established a diverse bacterial culture collection isolated from the rhizoplane of winter wheat (*Triticum aestivum* cv. Sheriff). Based on previously published data from Zervas *et al*. (2022) and Guan *et al*. (2024), we included some of the most abundant genera as well as rare genera found at the rhizoplane of the Sheriff cultivar at two different growth stages (Table 1).

**Table 1.**
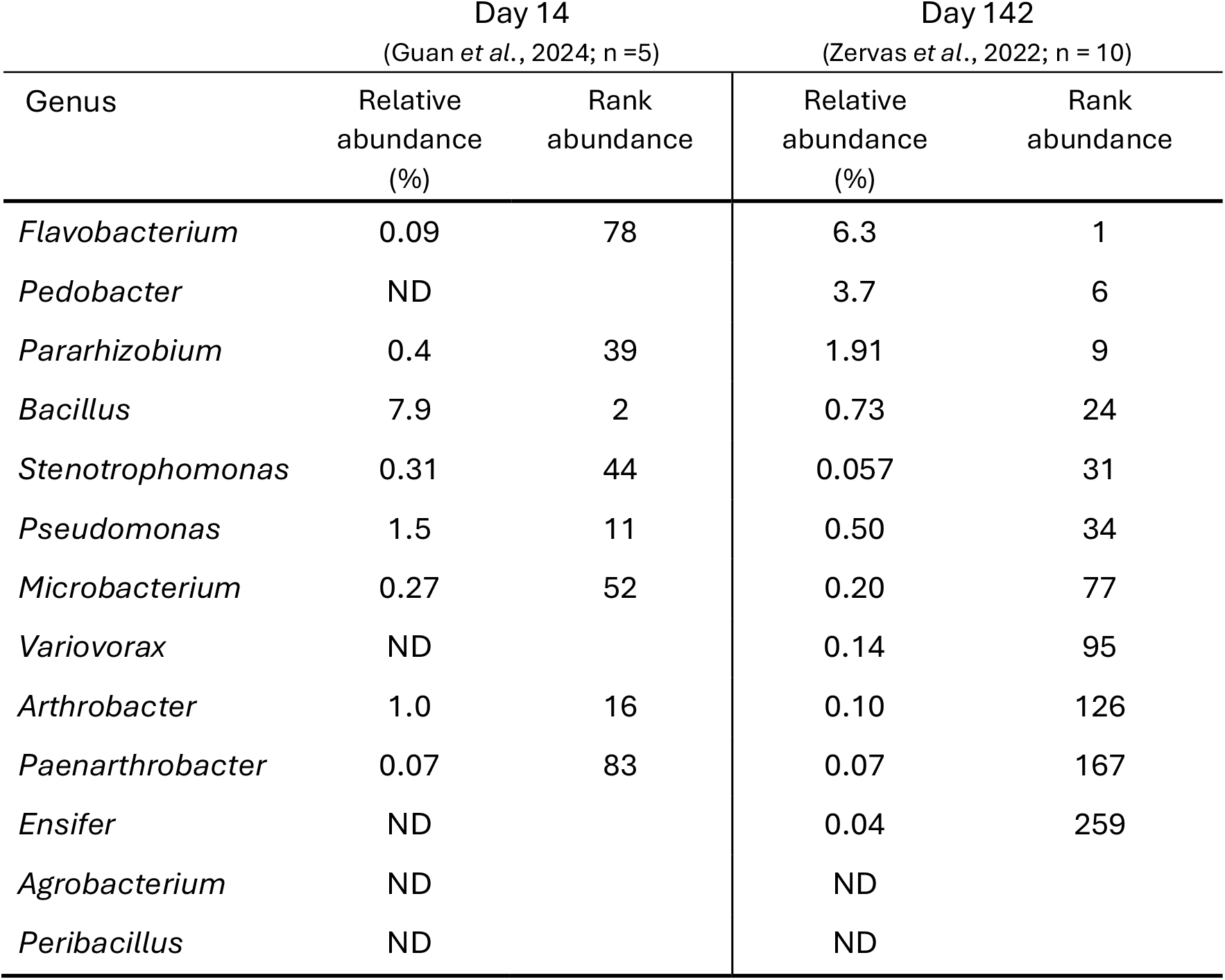
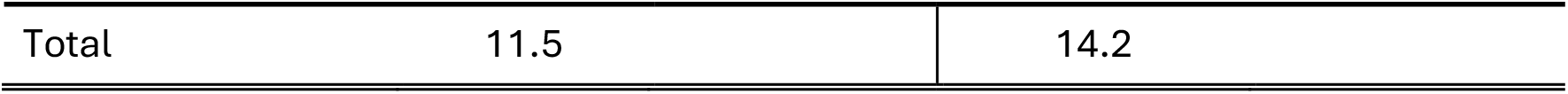
Mean relative abundance of SynCom genera detected by 16S rRNA gene sequencing in two previous studies. Rank abundance of the SynCom genera found in the total community. Days indicate days after sowing. ND: Not detected.

| Genus | Day 14<br>(Guan *et al.*, 2024; n =5) |  | Day 142<br>(Zervas *et al.*, 2022; n = 10) |  |
| --- | --- | --- | --- | --- |
|  | Relative abundance (%) | Rank abundance | Relative abundance (%) | Rank abundance |
| *Flavobacterium* | 0.09 | 78 | 6.3 | 1 |
| *Pedobacter* | ND |  | 3.7 | 6 |
| *Pararhizobium* | 0.4 | 39 | 1.91 | 9 |
| *Bacillus* | 7.9 | 2 | 0.73 | 24 |
| *Stenotrophomonas* | 0.31 | 44 | 0.057 | 31 |
| *Pseudomonas* | 1.5 | 11 | 0.50 | 34 |
| *Microbacterium* | 0.27 | 52 | 0.20 | 77 |
| *Variovorax* | ND |  | 0.14 | 95 |
| *Arthrobacter* | 1.0 | 16 | 0.10 | 126 |
| *Paenarthrobacter* | 0.07 | 83 | 0.07 | 167 |
| *Ensifer* | ND |  | 0.04 | 259 |
| *Agrobacterium* | ND |  | ND |  |
| *Peribacillus* | ND |  | ND |  |

|  |  |  |
| --- | --- | --- |
| Total | 11.5 | 14.2 |

Additionally, two genera, *Pseudomonas* and *Arthrobacter*, classified as core members of the wheat rhizosphere were selected (Simonin *et al*., 2020), as well as *Variovorax* due to its importance for root growth in *A. thaliana* (Finkel *et al*., 2020).

Ultimately, the SynCom comprised 13 strains from 13 genera: *Bacillus, Peribacillus* (Firmicutes), *Flavobacterium, Pedobacter* (Bacteroidota), *Arthrobacter, Microbacterium, Paenarthrobacter* (Actinobacteriota), *Agrobacterium, Ensifer, Pararhizobium, Pseudomonas, Stenotrophomonas* and *Variovorax* (Proteobacteria).

The 13 SynCom genera accounted for 12% and 14% of the genera detected in the rhizoplane community after 14 and 142 days of growth, respectively (Table 1). The selected strains were tested for various phenotypic traits to ensure a diverse functional potential of relevance for plant-microbe interactions (Table S1, Fig. S3-5).

### SynCom inoculation and plant development

The stable establishment and succession of the SynCom members on the wheat rhizoplane is crucial for further inferences on microbial community development and function, and derived effects (beneficial or detrimental) on plant health and performance. Here, we used gamma-irradiated soil, inoculated with the SynCom in the top soil, to strongly reduce the abundance of indigenous soil microorganisms, which would otherwise impede identification of the SynCom members during experiments.

Plants inoculated with the SynCom developed in a uniform way across replicates, for both shoot and root length (Fig. 1), and no indications of nutrient limitations or pathogen attack was observed during the four weeks of growth. The shoot height and root length reached ∼40 cm and ∼17.5 cm after 28 days of growth, respectively. We found no differences in shoot or root length between SynCom and NatCom nor were there any visual differences on plant growth and health between the inoculations (Fig. S6).

**Figure 1.**
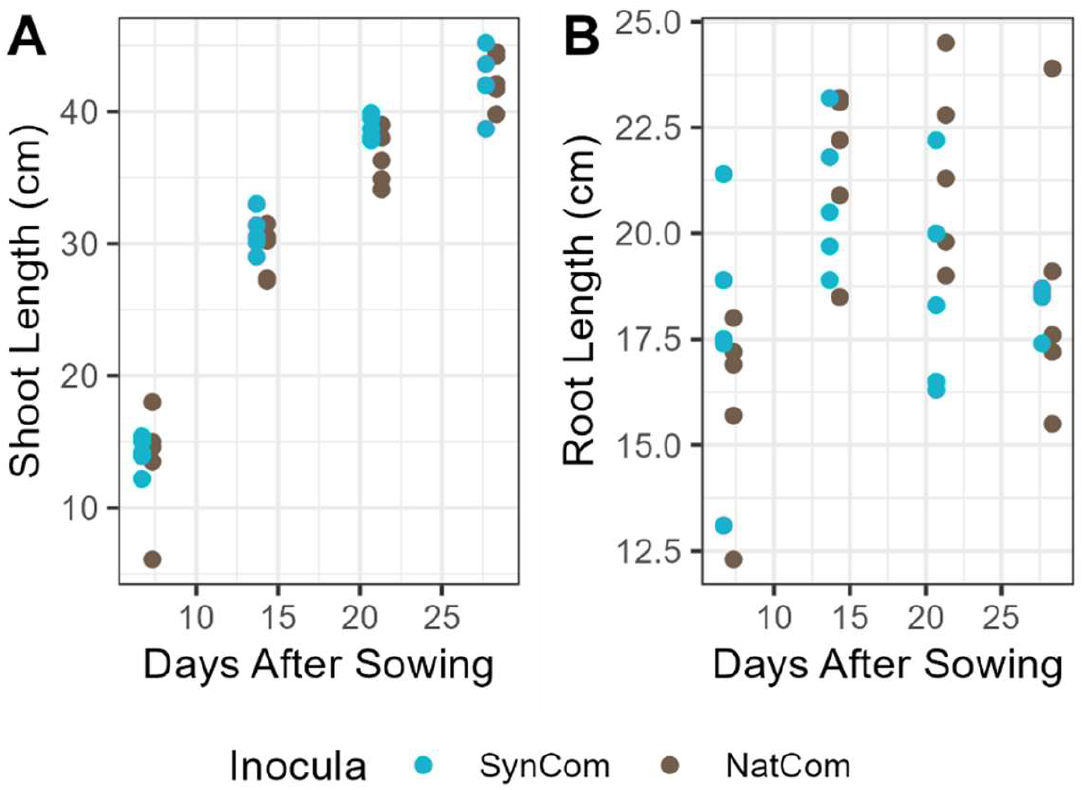
Shoot (A) and root (B) length at the different sampling times. Each point represents one plant.

### SynCom strains successfully colonized the roots of wheat

Based on 16S rRNA gene amplicon sequencing all SynCom strains successfully colonized the rhizoplane at Day 7 and all strains were, with few exceptions, part of the rhizoplane community at all sampling times (Fig. 2, Fig. S7).

**Figure 2.**
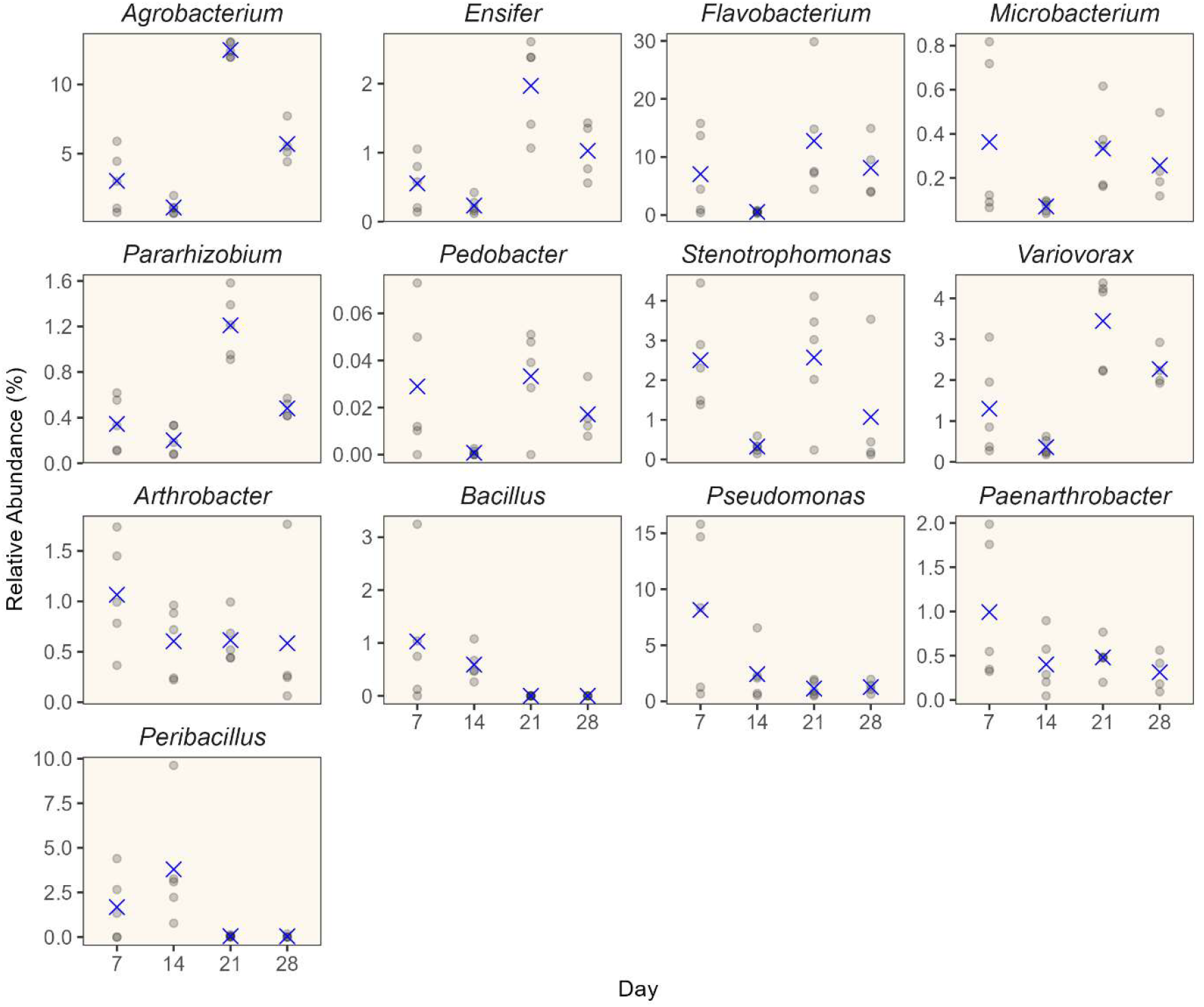
Relative abundance of SynCom genera in the gamma-irradiated soil and on wheat plants where SynCom was inoculated. The relative abundance of each genus is the sum of ASVs from each genus, found in starting SynCom inoculum (n = 5 plants). The relative abundances of each replicate plant are shown as shaded circles and the mean is shown as a cross.

We determined the proportion of each of the 13 SynCom genera originating from the starting inoculum or another source (gamma-irradiated soil or seed) based on ASVs (Fig. 3). For most of the genera, the majority of the ASVs came from the SynCom inoculum. The exceptions were *Bacillus* and *Pedobacter. Bacillus* was primarily consisting of non-SynCom ASVs at all four sampling time points, while *Pedobacter* showed a high proportion of ASVs not originating from the SynCom at Day 21 and 28. None of the ASVs in the starting SynCom, except for one ASV (*Peribacillus simplex* ASV13), were found in the gamma irradiated soil prior to inoculation, highlighting the applicability of the gamma-irradiated soil. In summary, this demonstrates that the added SynCom strains actively colonized the roots.

**Figure 3.**
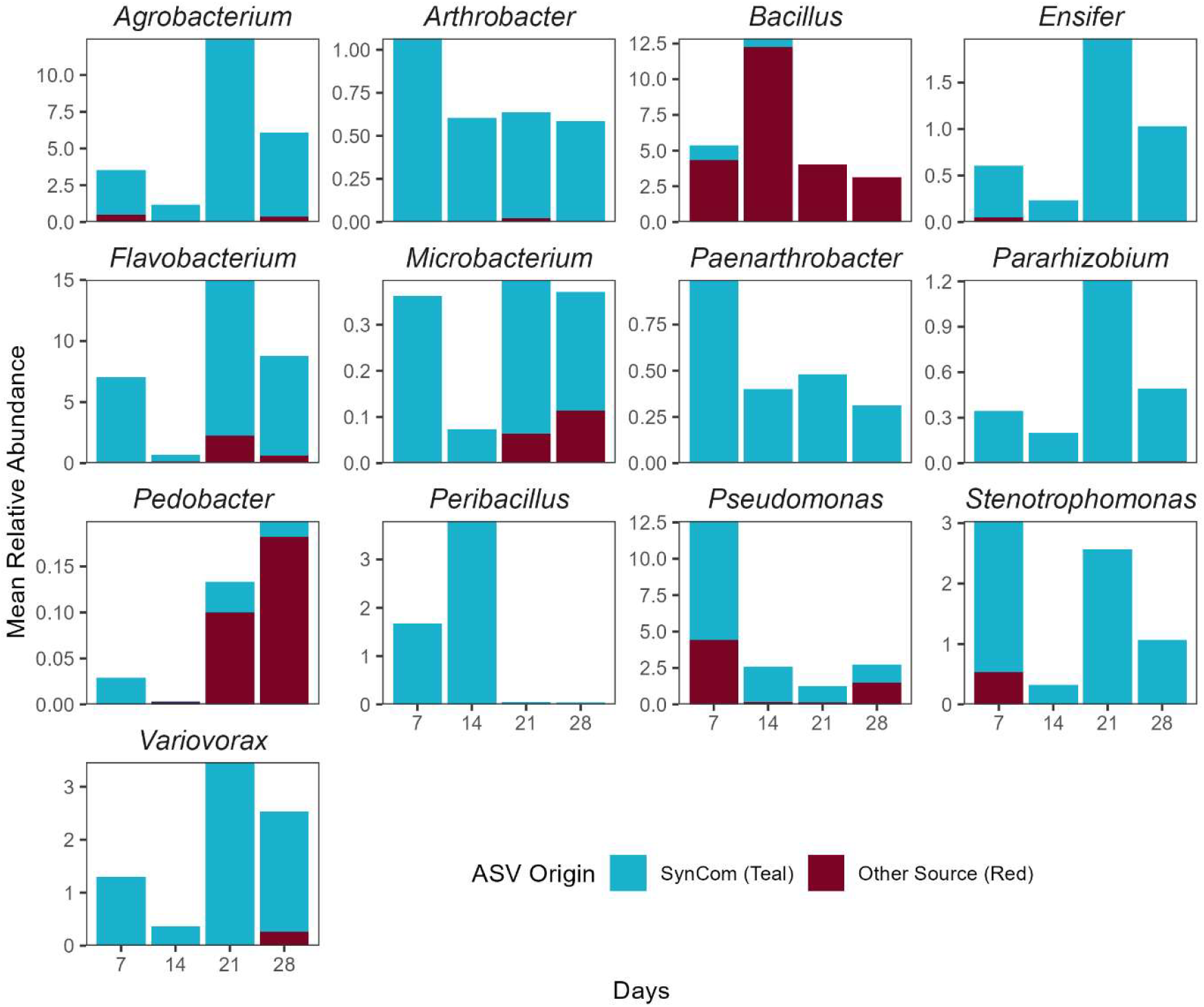
Origin of SynCom ASVs detected in the rhizoplane samples. The bars indicate the mean relative abundance (per day) of the total community of each genus encompassing all ASVs affiliated with the specific genus. *Peribacillus simplex* ASV13, found in both SynCom and gamma-irradiated soil is grouped under SynCom ASVs (teal). Other sources include soil or seed (red).

### The SynCom community decreases in variability over time

To use the SynCom for making inferences of what would happen on field scale, a robust model system is required with low between-replicate variability, and the development should resemble the community development in a natural community. The alpha diversity (Shannon index) determined based on all ASVs (i.e. including ASVs of seed and soil origin) increased over time in the SynCom inoculated plants (Fig. S8). Despite the initial difference in diversity between SynCom and NatCom inocula, no difference in alpha diversity between NatCom and SynCom treated plants was observed at any of the sampling days, although the variation among replicates was higher for NatCom inoculated samples (Fig. S8). Initial root community assembly is governed by stochastic processes (Dini-Andreote and Raaijmakers, 2018), followed by more deterministic processes. We therefore hypothesized that our SynCom would follow this pattern. To test this, we used the Bray Curtis dissimilarity of the entire rhizoplane community, a beta diversity index, as a metric for dissimilarity among communities at the four sampling times. We would expect the highest Bray Curtis dissimilarity at Day 7 due to the initial stochastic assembly. Indeed, at Day 7, the SynCom and NatCom communities were highly dissimilar among replicates (Fig. 4A), suggesting more randomness in community assembly at this time point. At Day 14, the dissimilarity among replicates differed between SynCom and NatCom, whereafter communities from both treatments stabilized. This stabilization of the Bray Curtis dissimilarity supporting that community assembly is less stochastic at later development stages.

**Figure 4.**
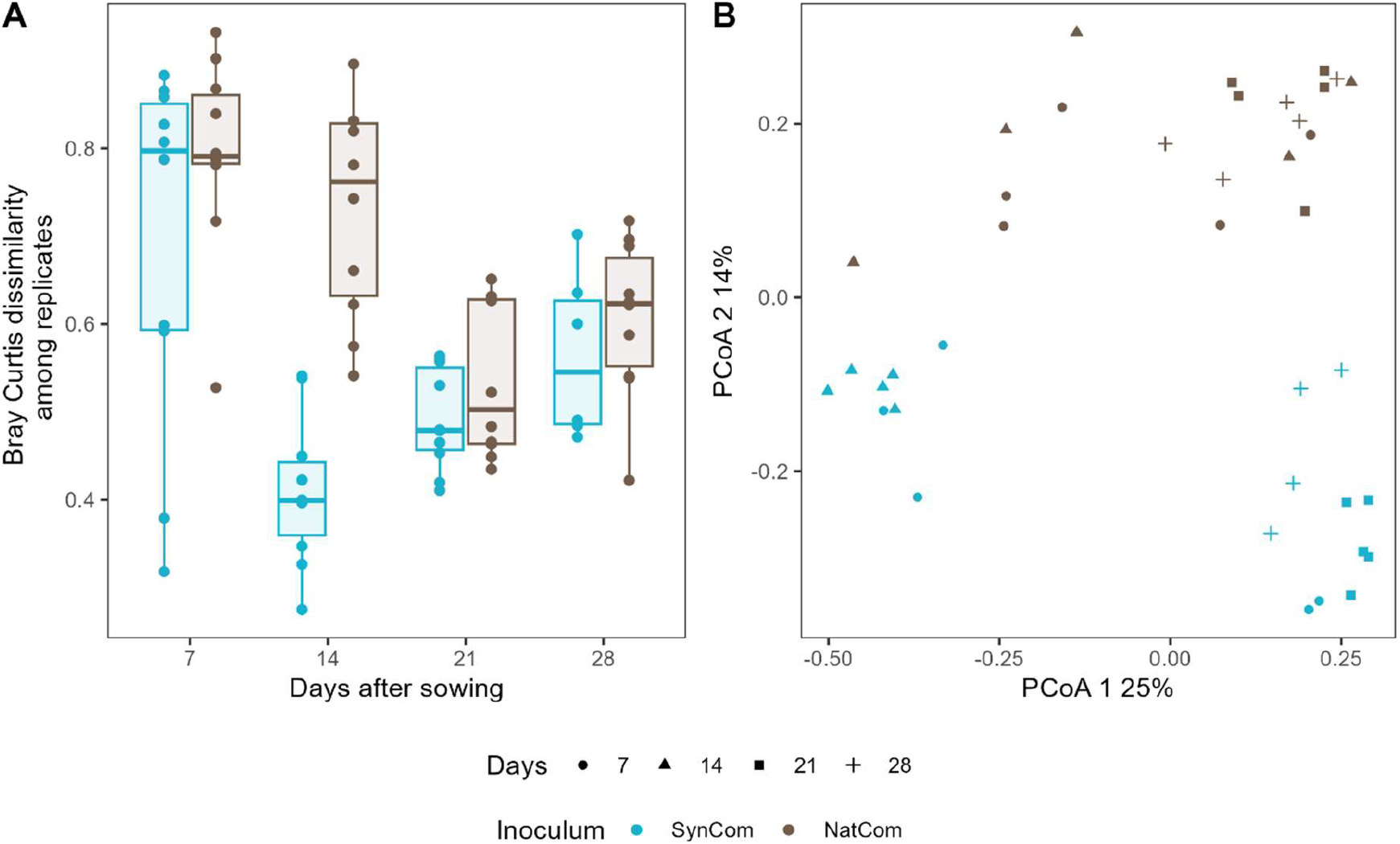
(A) Bray-Curtis dissimilarities of 16S rRNA amplicons among replicates of SynCom or NatCom communities at the four sampling days. Each point represents a comparison. The boxplot shows the median (horizontal line), the interquartile range (25th-75th percentile), and the whiskers extend from the hinges to the largest value no further than 1.5x the interquartile range. (B) PCoA of Bray-Curtis dissimilarities.

The PCoA ordination showed a separation between SynCom communities from Day 14, and those from Day 21 and 28 samples (Fig. 4B), which could be linked to the reduction in relative abundance of *Massilia* between Day 14 and 21 (Fig. S9). While *Massilia* was not found in the soil, we isolated seed endophytic *Massilia* (Table S3). Overall, our data show that despite a lower alpha diversity of the inoculated SynCom compared to the NatCom (Fig. S8), the diversity measures of the SynCom community followed the same overall trends as the NatCom rhizoplane communities. This supports the hypothesis that initial root microbiome assembly is stochastic followed by a non-random assembly. Furthermore, the data indicate that seed endophytes comprise a large proportion of the rhizoplane microbiome during early plant development, before SynCom members are recruited and colonize the roots.

We compared the assembly and succession of the SynCom community with the NatCom community development by tracking all NatCom ASVs belonging to the 13 SynCom genera over time (Fig. 5). Two genera, *Bacillus* and *Peribacillus* peaked in relative abundance at Day 14. Most other genera had the lowest relative abundance at Day 14, followed by an increase in relative abundance to Day 21. The genera *Bacillus, Flavobacterium, Microbacterium, Pararhizobium, Pseudomonas, Agrobacterium and Pedobacter* accounted for similar amounts of the total abundance in both communities in the later time points (Fig. 5), despite the differences in relative abundance in the starting community (Day 0) (Fig. S11). *Arthrobacter* and *Paenarthrobacter* contributed more to the total community at the first three time points in the SynCom compared to the NatCom, while being comparable at Day 28. These results suggest that the selected genera can colonize a specific proportion of the rhizoplane irrespective of starting concentration.

**Figure 5.**
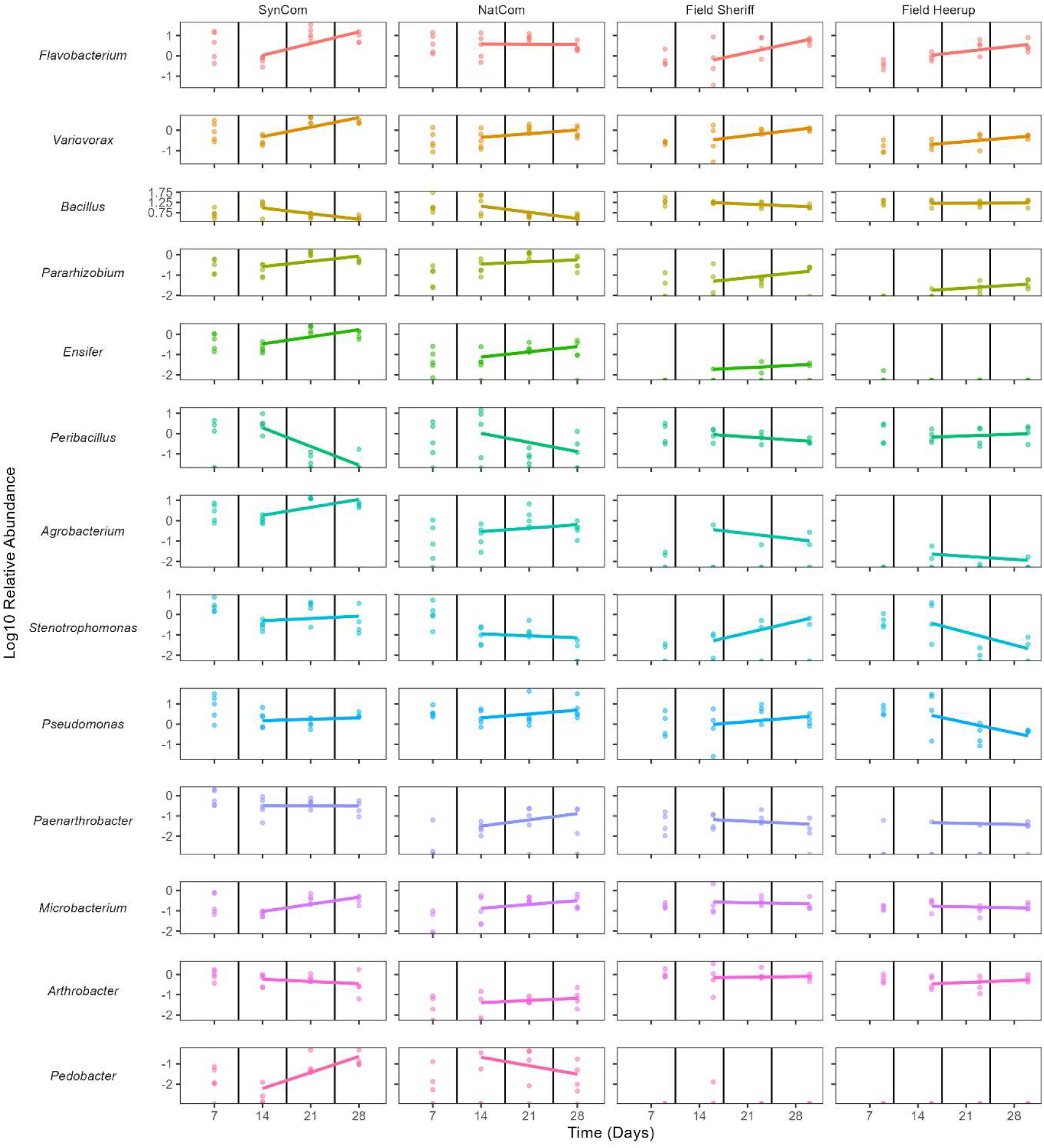
Log10-normalized read counts of all ASVs assigned to the taxonomy of SynCom genera in SynCom, NatCom and field samples. The inserted regression lines show the ANCOVA models fitted within genera from Day 14 to Day 28. The genera are ordered according to statistical evidence for the same sign of the relative abundance for SynCom and Field Sheriff samples (see Table 3 below) using the COST procedure.

The temporal dynamics of the relative abundance of each SynCom member was compared to the corresponding genera in the NatCom with a new statistical method, COST *(Conservative One-Sided Test)*. There was evidence (marginal p-values below 0.05) for the same direction of the temporal dynamics for four genera; *Bacillus* and *Peribacillus* were decreasing, and *Microbacterium* and *Variovorax* were increasing (Table 2). A similar COST procedure can be used to test for different signs using the null hypothesis of equal signs. There was no evidence for different direction of temporal dynamics for any of the genera.

**Table 2.**
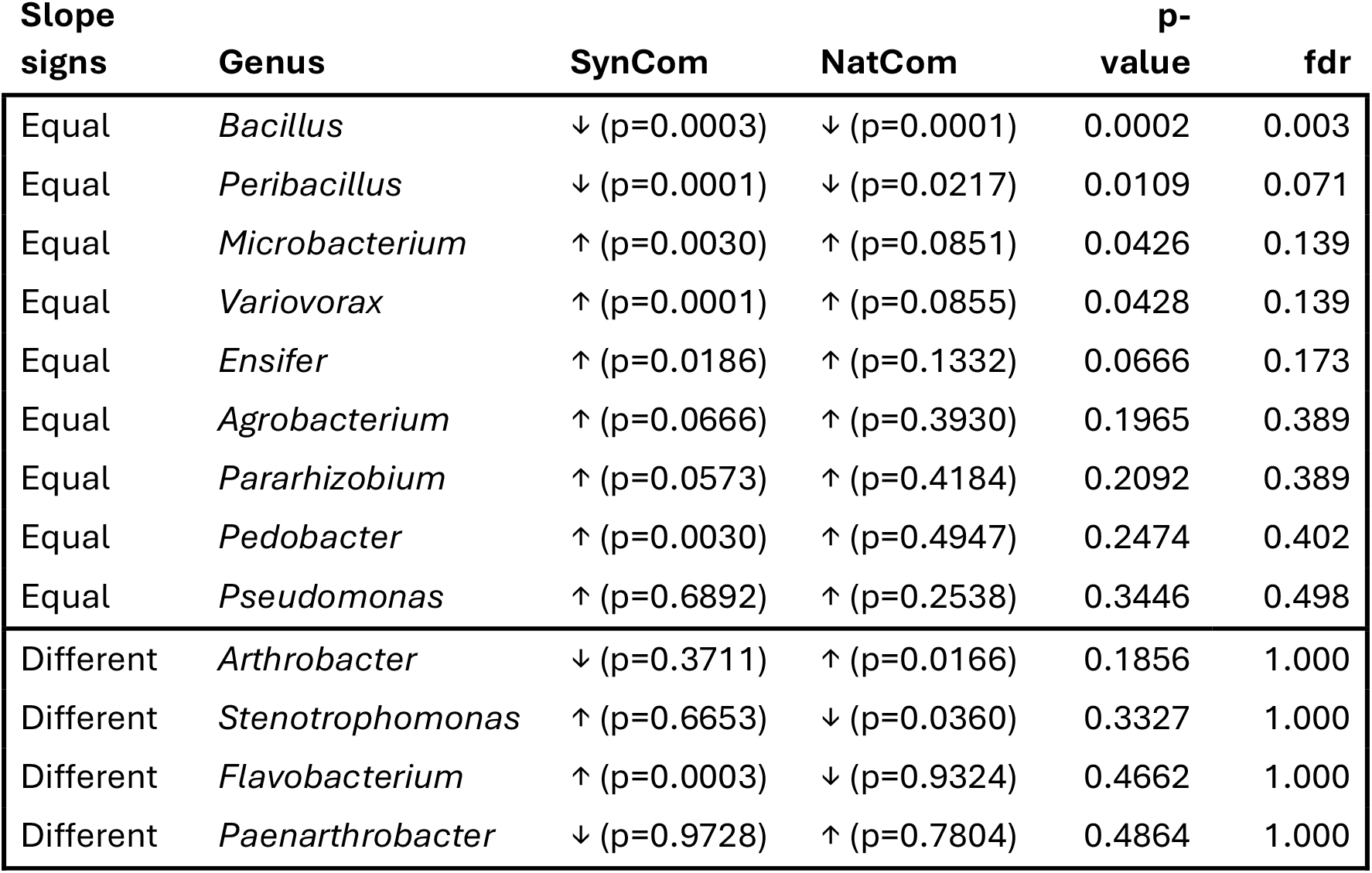
Comparison of the sign of the growth rates in SynCom and NatCom. The third and fourth columns specify if the relative abundances are increasing (↑) or decreasing (↓) in SynCom and NatCom, respectively, with p-value for the marginal two-sided T-tests stated in the parentheses. The fifth and sixth column specify p-values and associated false discovery rates for scientific hypothesis of equal signs (first 9 rows) and different signs (last 4 rows). Genera within equal and different signs are ordered according to increasing p-values.

| Slope signs | Genus | SynCom | NatCom | p-value | fdr |
| --- | --- | --- | --- | --- | --- |
| Equal | *Bacillus* | ↓ (p=0.0003) | ↓ (p=0.0001) | 0.0002 | 0.003 |
| Equal | *Peribacillus* | ↓ (p=0.0001) | ↓ (p=0.0217) | 0.0109 | 0.071 |
| Equal | *Microbacterium* | ↑ (p=0.0030) | ↑ (p=0.0851) | 0.0426 | 0.139 |
| Equal | *Variovorax* | ↑ (p=0.0001) | ↑ (p=0.0855) | 0.0428 | 0.139 |
| Equal | *Ensifer* | ↑ (p=0.0186) | ↑ (p=0.1332) | 0.0666 | 0.173 |
| Equal | *Agrobacterium* | ↑ (p=0.0666) | ↑ (p=0.3930) | 0.1965 | 0.389 |
| Equal | *Pararhizobium* | ↑ (p=0.0573) | ↑ (p=0.4184) | 0.2092 | 0.389 |
| Equal | *Pedobacter* | ↑ (p=0.0030) | ↑ (p=0.4947) | 0.2474 | 0.402 |
| Equal | *Pseudomonas* | ↑ (p=0.6892) | ↑ (p=0.2538) | 0.3446 | 0.498 |
| Different | *Arthrobacter* | ↓ (p=0.3711) | ↑ (p=0.0166) | 0.1856 | 1.000 |
| Different | *Stenotrophomonas* | ↑ (p=0.6653) | ↓ (p=0.0360) | 0.3327 | 1.000 |
| Different | *Flavobacterium* | ↑ (p=0.0003) | ↓ (p=0.9324) | 0.4662 | 1.000 |
| Different | *Paenarthrobacter* | ↓ (p=0.9728) | ↑ (p=0.7804) | 0.4864 | 1.000 |

**Table 3.** Comparison of the sign of the growth rates in SynCom and Field Sheriff. Evidence for equal (first 10 rows) and different (last 2 rows) signs was evaluated similarly as described in Table 2. *Pedobactor* was not included due to the very low abundance for Field Sheriff.

| Slope signs | Genus | SynCom | Field Sheriff | p-value | fdr |
| --- | --- | --- | --- | --- | --- |
| Equal | *Flavobacterium* | ↑ ($p=0.0003$) | ↑ ($p=0.0019$) | 0.0010 | 0.012 |
| Equal | *Variovorax* | ↑ ($p=0.0001$) | ↑ ($p=0.0184$) | 0.0092 | 0.055 |
| Equal | *Bacillus* | ↓ ($p=0.0003$) | ↓ ($p=0.1730$) | 0.0865 | 0.269 |
| Equal | *Pararhizobium* | ↑ ($p=0.0573$) | ↑ ($p=0.1794$) | 0.0897 | 0.269 |
| Equal | *Ensifer* | ↑ ($p=0.0202$) | ↑ ($p=0.3370$) | 0.1685 | 0.404 |
| Equal | *Peribacillus* | ↓ ($p=0.0001$) | ↓ ($p=0.4535$) | 0.2268 | 0.441 |
| Equal | *Agrobacterium* | ↑ ($p=0.0666$) | ↑ ($p=0.5146$) | 0.2573 | 0.441 |
| Equal | *Stenotrophomonas* | ↑ ($p=0.6653$) | ↑ ($p=0.6707$) | 0.3354 | 0.459 |
| Equal | *Pseudomonas* | ↑ (p=0.6892) | ↑ (p=0.2925) | 0.3446 | 0.459 |
| Equal | *Paenarthrobacter* | ↓ (p=0.9728) | ↓ (p=0.2012) | 0.4864 | 0.584 |
| Different | *Microbacterium* | ↑ (p=0.0030) | ↓ (p=0.7231) | 0.3616 | 1.000 |
| Different | *Arthrobacter* | ↓ (p=0.3711) | ↑ (p=0.8315) | 0.4158 | 1.000 |

### Temporal dynamics show similar patterns in the field, but the trend is cultivar dependent

To determine whether the assembly of the SynCom under controlled conditions would reflect assembly of microbial communities in the field, and whether it was cultivar dependent, we collected rhizoplane samples from the field of two winter wheat cultivars (*cv*. Sheriff, used for SynCom and NatCom experiments, and *cv*. Heerup) over a 4-week period in the fall of 2023. Seeds were sown without surface sterilization or pre-germination, in contrast to our SynCom and NatCom setup, to mimic agricultural practices. Expectedly, the alpha diversity was higher in the field soil than in the SynCom or NatCom inocula (Fig. S8). This was also true for the alpha diversity for rhizoplane communities of both cultivars grown in the field at all sampling time points (Fig. S8). The day of sampling explained 9% of the variation in the data (PERMANOVA, p < 0.001), and was thus more important than cultivar (PERMANOVA, R^2^ = 0.06, p = 0.002). The most dominant genera in the field were *Bacillus, Paenibacillus* and *Massilia* across the sampling period for both cultivars (Fig. S12). Except for *Agrobacterium, Ensifer* and *Pedobacter*, the rest of the SynCom genera were also detected in the field soil prior to sowing (Fig. S11). *Ensifer* and *Pedobacter* were also very low abundant or absent on the rhizoplane of both cultivars (Fig. 5). Despite this, the total relative abundance of SynCom genera comprised comparable proportions of the total communities regardless of experimental design.

When comparing slopes, we found that the majority of the SynCom members had slopes of equal signs when compared to Field grown plants (Table 3). There was evidence (marginal p-values < 0.05) for the same direction of the temporal dynamics for two genera; *Flavobacterium* and *Variovorax* that were both increasing. There was no evidence for different direction of temporal dynamics for any of the genera.

For Heerup, a winter wheat variety with a different microbiome (Zervas *et al*., 2022; Herms *et al*., 2025), there was evidence (marginal p-values < 0.05) for the same direction of the temporal dynamics for three genera; *Pararhizobium, Flavobacterium*, and *Variovorax* (Table 4). Notably these three genera also have the same direction for Field Sheriff. For the remaining there were different directions between SynCom and Field Heerup, but none of these were statistically significant.

**Table 4.**
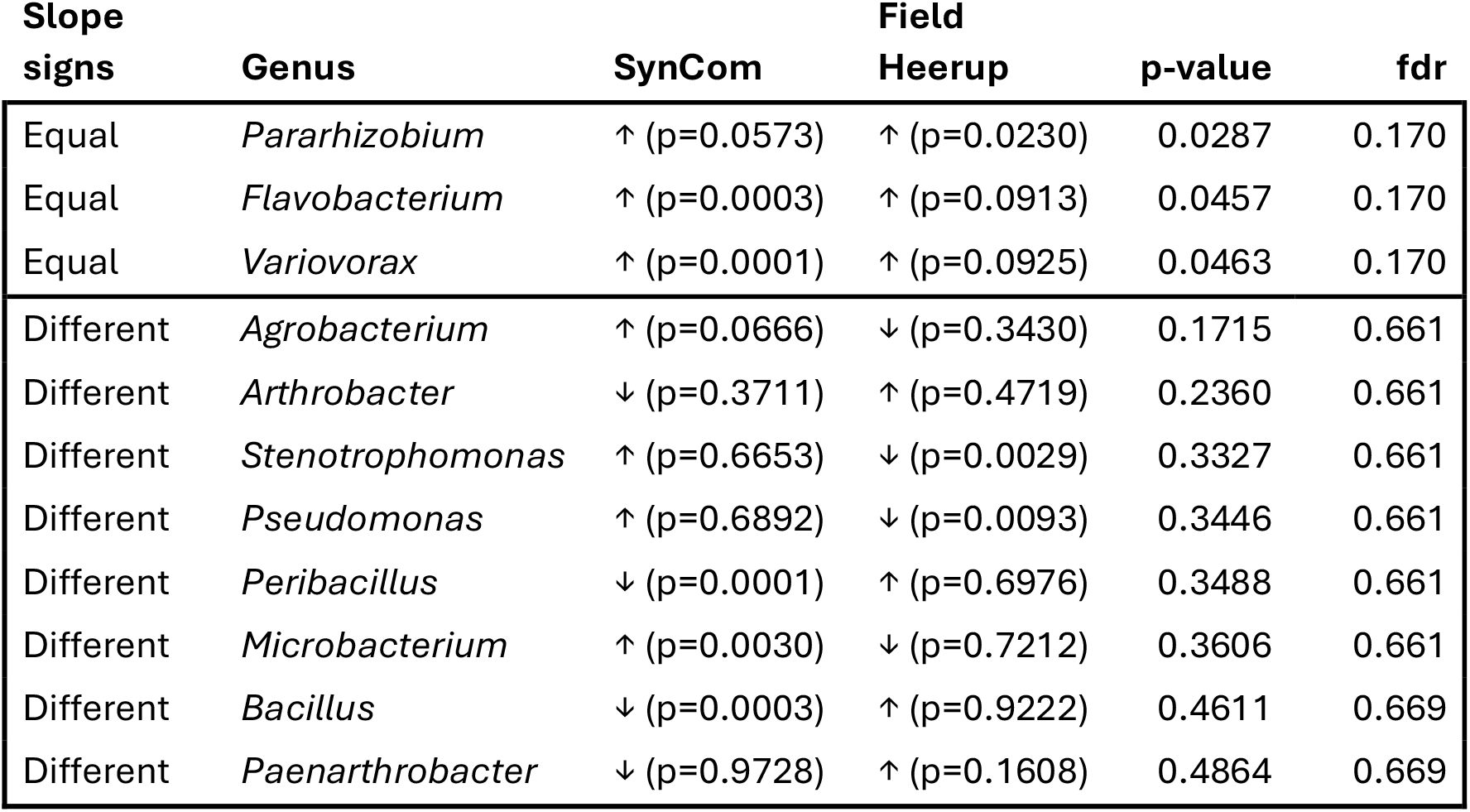
Comparison of the sign of the growth rates in SynCom and Field Sheriff. Evidence for equal (first 3 rows) and different (last 8 rows) signs was evaluated similarly as described in Table 2. *Ensifer* and *Pedobactor* were not included due to the very low abundance for Field Heerup.

## Discussion

Here, we showed that a 13-member bacterial SynCom originating from the roots of winter wheat *cv*. Sheriff colonized the roots after inoculation in gamma-irradiated soil in a similar manner as observed for field grown Sheriff plants. By applying the SynCom to the soil instead of the seed we can study active rhizosphere colonization, contrasting previous studies (Niu *et al*., 2017; Simonin *et al*., 2023). Another advantage is the use of gamma-irradiated soil which mimics natural soil more than other substrates used in previous SynCom experiments (Ma *et al*., 2021; Zhou *et al*., 2022; Lin *et al*., 2024) and thus allows to test rhizosphere community assembly under more realistic conditions. The approaches taken in previous studies are likely explanations for the difficulties often experienced when transferring knowledge from lab studies to the field (Sessitsch *et al*., 2019).

These findings underpin the strong selective forces that drive rhizoplane community development, at least in the first month after sowing. Further, it shows that the selected SynCom can be used to recreate the overall structures of a natural community showcasing the great potential for SynComs to shed light on plant-microbe interaction in future research.

We found no differences in shoot biomass and length between the SynCom and NatCom inoculated plants suggesting that the SynCom provides the necessary functions for plant growth development in the early stages. This complements the findings from maize, radish and tomato that have shown that 7-15 SynCom members are sufficient for early plant development (Niu *et al*., 2017; Schmitz *et al*., 2022; Simonin *et al*., 2023).

All SynCom strains colonized the rhizoplane over the four weeks of growth. Only one of the SynCom ASVs (*Peribacillus simplex* ASV13) was found in the soil prior to inoculation, and hence, we conclude that the SynCom ASVs found on the roots originated from the SynCom inoculum. None of the strains completely dominated the rhizoplane indicative of a niche preference for each SynCom member. Previous studies using single strains in gnotobiotic systems with wheat, barley and maize have shown that there is a maximum colonization potential, even without competition from other than seed endophytes (Bennett and Lynch, 1981). In line with this, Guan *et al*. (2024) showed that inoculation with *Pseudomonas fluorescens* SBW25 did not change the relative abundance of *Pseudomonas* on wheat roots compared to untreated seeds . Taken together, this suggests that the plants recruit a restricted number of bacteria from a given genus due to environmental conditions or limitation of space on the roots. This should be considered when applying bioinoculants as the main competitor might be species from the same genus due to niche preferences.

Despite surface sterilization and the use of irradiated soil, we were not able to remove all other sources of bacteria. This was especially apparent for *Bacillus*, as the majority of the ASVs from this genus was from other sources than the SynCom. It is a known seed endophyte (Díaz Herrera *et al*., 2016), but was also present in the gamma-irradiated soil. Hence, a higher dose of gamma-irradiation is required to remove dormant *Bacillus* from the soil, if this genus is of particular interest in the SynCom. In addition, the high proportion of *Massilia* on the SynCom treated wheat roots in the initial 14 days can be ascribed to its presence as a seed endophyte (Table S3) in multiple plant species, e.g. pea, maize, sorghum and wheat (Johnston-Monje *et al*., 2021). This is consistent with the NatCom treated plants and the field-grown plants, as well as germinating wheat seeds (Bornø *et al*., *submitted*), wheat roots (Guan *et al*., 2024) and cucumber (Ofek *et al*., 2012) during early plant development.

In support of our first hypothesis, we observed the largest variation in community composition (beta diversity) among replicates after 7 days for the SynCom communities. For NatCom the variation in community composition was evenly high at Day 7 and 14. The difference at Day 14 probably reflects that the SynCom consisted of strains isolated from the wheat rhizosphere and were added as actively growing cells, and thus could colonize faster than the bacteria from the NatCom. Alternatively, the lower diversity of the SynCom could have resulted in a faster equilibrium among species and a higher root colonization of seed borne bacteria compared to the NatCom. In fact, the high relative abundance of the *Massilia* found at Day 7 could support this. The high variation in beta diversity during early stages of plant growth was also shown for maize (Rüger *et al*., 2021; Xiong *et al*., 2021). The lower beta diversity among replicates at later developmental stages for the SynCom and NatCom supports a previously proposed model with stochastic processes being more important in early stages of development, while deterministic processes become more dominant over time (Dini-Andreote and Raaijmakers, 2018).

As growth rate potential is conserved at the genus level (Weissman *et al*., 2021), we hypothesized that a single strain could be representative of the growth pattern/temporal dynamics. Overall, the temporal dynamics of 10 strains/genera in the SynCom, except *Pedobacter, Arthrobacter and Microbacterium* support this hypothesis, as they resembled those of the field grown Sheriff cultivar. This is despite the lower diversity in the SynCom inoculum and lack of archaea and eukaryotes compared to the field soil and the differences in seed treatment and lower temperatures for the field grown plants.

In contrast to the high similarity in dynamics of the 13 SynCom genera for Sheriff across systems, fewer similarities were found for the Heerup cultivar. This could be explained by wheat-cultivar specific recruitment of members within a genus, as shown for *Pseudomonas* in the two wheat cultivars (Herms *et al*., 2025). This provides one explanation why translating results from lab to field often fails, when even temporal dynamics of genera are different between two cultivars.

Since the conducted growth chamber experiment only lasted for 28 days, further experiments are warranted for testing long-term related hypotheses. In fact, this is a general challenge in the research field (Martins *et al*., 2023). Another aspect is the observations of zero abundance of some genera that can be due to either unsuccessful colonization or too low abundance to be detected. Since bacterial abundance is best modelled on a logarithmic scale these zeros pose a challenge. An often-used solution is to add a small positive number to all observations. However, this solution is *ad hoc*, and results may change with the number used. Here, we chose to interpret a zero as the observation that this abundance is lower than all the non-zero abundances within that genus. This solution still entails a modelling choice and hence impacts the precise p-values in the statistical computations. However, we believe that this is less *ad hoc* than adding an arbitrary small number. An alternative is to only investigate genera without zero observations, but this would exclude several genera (2 for SynCom, 6 for NatCom, 5 for Field Sheriff, and 4 for Field Heerup).

In conclusion, we demonstrated the potential of applying a SynCom to study rhizoplane microbiome assembly on wheat in soil. Its members actively colonized the wheat roots and despite lacking eukaryotes and archaea, it developed in accordance with field communities of the same wheat cultivar. Overall, this supports the notion that there is a certain carrying capacity for each community on the roots due to niche preferences. Recognizing this will be helpful in advancing efforts on the application of beneficial microbes to seeds for root colonization.

Our approach with inoculating the SynCom into gamma-irradiated soil provides a promising framework as a more realistic scenario, compared to seed inoculation and the use of hydroponics to determine governmental factors of microbial community assembly at the sub-genus level and to leverage studies of plant-microbe and microbe-microbe interactions at the root-soil interface. Thus, we believe that this approach will help bridge the gap between lab and field.

## Data availability

Raw 16S rRNA gene amplicon sequences have been deposited in the NCBI Sequence Read Archive (SRA) database under Bioproject accession number PRJNA1189047. The scripts describing data treatment and the COST procedure are available at: https://github.com/BakDK/Wheat-SynCom

## Acknowledgements

We thank Dorthe Thybo Ganzhorn for assistance with sowing and watering the plants used for cultivation. We thank Dorette Müller-Stöver for providing the gamma irradiated soil.

Thank you for the help with field sampling to our research group members: Xingyun Yi, Sarah Buch Nielsen, Benjamin Nils Thorn, Jabeen Ahmad, Courtney Horn Herms, Dorthe Thybo Ganzhorn and Klara Gunnarsen. This work was conducted within the INTERACT project funded by Novo Nordisk Foundation (grant no. NNF19SA0059360). Frederik Bak was supported by Independent Research Fund Denmark (grant no. 10.46540/2031-00010B).

## References

Beckers, B., Op De Beeck, M., Thijs, S., Truyens, S., Weyens, N., Boerjan, W., and Vangronsveld, J. (2016) Performance of 16s rDNA Primer Pairs in the Study of Rhizosphere and Endosphere Bacterial Microbiomes in Metabarcoding Studies. Frontiers in Microbiology 7:.

Bennett, R.A. and Lynch, J.M. (1981) Colonization potential of bacteria in the rhizosphere. Current Microbiology 6: 137–138.

van der Bom, F., Magid, J., and Jensen, L.S. (2017) Long-term P and K fertilisation strategies and balances affect soil availability indices, crop yield depression risk and N use. European Journal of Agronomy 86: 12–23.

Callahan, B.J., McMurdie, P.J., Rosen, M.J., Han, A.W., Johnson, A.J.A., and Holmes, S.P. (2016) DADA2: High-resolution sample inference from Illumina amplicon data. Nature Methods 13: 581–583.

Carlström, C.I., Field, C.M., Bortfeld-Miller, M., Müller, B., Sunagawa, S., and Vorholt, J.A. (2019) Synthetic microbiota reveal priority effects and keystone strains in the Arabidopsis phyllosphere. Nature Ecology & Evolution 3: 1445–1454.

Compant, S., Samad, A., Faist, H., and Sessitsch, A. (2019) A review on the plant microbiome: Ecology, functions, and emerging trends in microbial application. Journal of Advanced Research 19: 29–37.

Díaz Herrera, S., Grossi, C., Zawoznik, M., and Groppa, M.D. (2016) Wheat seeds harbour bacterial endophytes with potential as plant growth promoters and biocontrol agents of <i>Fusarium graminearum<\i>. Microbiological Research 186–187: 37–43.

Dini-Andreote, F. and Raaijmakers, J.M. (2018) Embracing Community Ecology in Plant Microbiome Research. Trends in Plant Science 23: 467–469.

Finkel, O.M., Salas-González, I., Castrillo, G., Conway, J.M., Law, T.F., Teixeira, P.J.P.L., et al. (2020) A single bacterial genus maintains root growth in a complex microbiome. Nature 587: 103–108.

Goslee, S.C. and Urban, D.L. (2007) The ecodist Package for Dissimilarity-based Analysis of Ecological Data. Journal of Statistical Software 22: 1–19.

Guan, Y., Bak, F., Hennessy, R.C., Horn Herms, C., Elberg, C.L., Dresbøll, D.B., et al. (2024) The potential of Pseudomonas fluorescens SBW25 to produce viscosin enhances wheat root colonization and shapes root-associated microbial communities in a plant genotype-dependent manner in soil systems. mSphere 0: e00294–24.

Harbort, C.J., Hashimoto, M., Inoue, H., Niu, Y., Guan, R., Rombolà, A.D., et al. (2020) Root-Secreted Coumarins and the Microbiota Interact to Improve Iron Nutrition in Arabidopsis. Cell Host & Microbe 28: 825-837.e6.

Herms, C.H., Hennessy, R.C., Bak, F., Guan, Y., Browne, P.D., Nielsen, T.K., et al. (2024) Pseudomonas taxonomic and functional microdiversity in the wheat rhizosphere is cultivar-dependent and links to disease resistance profile and root diameter. 2024.06.13.598955.

Herms, C.H., Hennessy, R.C., Bak, F., Guan, Y., Browne, P.D., Nielsen, T.K., et al. (2025) Pseudomonas taxonomic and functional microdiversity in the wheat rhizosphere is cultivar-dependent and links to disease resistance profile and root diameter. Applied Soil Ecology 211: 106116.

Jing, J., Garbeva, P., Raaijmakers, J.M., and Medema, M.H. (2024) Strategies for tailoring functional microbial synthetic communities. The ISME Journal 18: wrae049.

Johnston-Monje, D., Gutiérrez, J.P., and Lopez-Lavalle, L.A.B. (2021) Seed-Transmitted Bacteria and Fungi Dominate Juvenile Plant Microbiomes. Frontiers in Microbiology 12:.

Lin, H.-H., Torres, M., Adams, C.A., Andeer, P.F., Owens, T.K., Zhalnina, K., et al. (2024) Impact of Inoculation Practices on Microbiota Assembly and Community Stability in a Fabricated Ecosystem. Phytobiomes Journal 8: 155–167.

López, J.L., Fourie, A., Poppeliers, S.W.M., Pappas, N., Sánchez-Gil, J.J., De Jonge, R., and Dutilh, B.E. (2023) Growth rate is a dominant factor predicting the rhizosphere effect. The ISME Journal 17: 1396–1405.

Ma, K.-W., Niu, Y., Jia, Y., Ordon, J., Copeland, C., Emonet, A., et al. (2021) Coordination of microbe–host homeostasis by crosstalk with plant innate immunity. Nat Plants 7: 814–825.

Martins, S.J., Pasche, J., Silva, H.A.O., Selten, G., Savastano, N., Abreu, L.M., et al. (2023) The Use of Synthetic Microbial Communities to Improve Plant Health. Phytopathology® 113: 1369–1379.

McMurdie, P.J. and Holmes, S. (2013) phyloseq: An R Package for Reproducible Interactive Analysis and Graphics of Microbiome Census Data. PLOS ONE 8: e61217.

Mendes, R., Garbeva, P., and Raaijmakers, J.M. (2013) The rhizosphere microbiome: significance of plant beneficial, plant pathogenic, and human pathogenic microorganisms. FEMS Microbiology Reviews 37: 634–663.

Niu, B., Paulson, J.N., Zheng, X., and Kolter, R. (2017) Simplified and representative bacterial community of maize roots. Proceedings of the National Academy of Sciences 114: E2450–E2459.

Northen, T.R., Kleiner, M., Torres, M., Kovács, Á.T., Nicolaisen, M.H., Krzyżanowska, D.M., et al. (2024) Community standards and future opportunities for synthetic communities in plant–microbiota research. Nat Microbiol 9: 2774–2784.

Ofek, M., Hadar, Y., and Minz, D. (2012) Ecology of Root Colonizing Massilia (Oxalobacteraceae). PLoS One 7: e40117.

Oksanen, J., Blanchet, F.G., Friendly, M., Kindt, R., Legendre, P., McGlinn, D., et al. (2020) vegan: Community Ecology Package.

Pfeilmeier, S., Werz, A., Ote, M., Bortfeld-Miller, M., Kirner, P., Keppler, A., et al. (2024) Leaf microbiome dysbiosis triggered by T2SS-dependent enzyme secretion from opportunistic Xanthomonas pathogens. Nat Microbiol 9: 136–149.

Quast, C., Pruesse, E., Yilmaz, P., Gerken, J., Schweer, T., Yarza, P., et al. (2013) The SILVA ribosomal RNA gene database project: improved data processing and web-based tools. Nucleic Acids Res 41: D590–D596.

R Core Team (2020) R: A language and environment for statistical computing.

Rüger, L., Feng, K., Dumack, K., Freudenthal, J., Chen, Y., Sun, R., et al. (2021) Assembly Patterns of the Rhizosphere Microbiome Along the Longitudinal Root Axis of Maize (Zea mays L.). Front Microbiol 12:.

Schloss, P.D. (2024) Rarefaction is currently the best approach to control for uneven sequencing effort in amplicon sequence analyses. mSphere 9: e00354–23.

Schmitz, L., Yan, Z., Schneijderberg, M., de Roij, M., Pijnenburg, R., Zheng, Q., et al. (2022) Synthetic bacterial community derived from a desert rhizosphere confers salt stress resilience to tomato in the presence of a soil microbiome. ISME J 16: 1907–1920.

Schuirmann, D.J. (1987) A comparison of the Two One-Sided Tests Procedure and the Power Approach for assessing the equivalence of average bioavailability.

Sessitsch, A., Pfaffenbichler, N., and Mitter, B. (2019) Microbiome Applications from Lab to Field: Facing Complexity. Trends in Plant Science 24: 194–198.

Simonin, M., Dasilva, C., Terzi, V., Ngonkeu, E.L.M., Diouf, D., Kane, A., et al. (2020) Influence of plant genotype and soil on the wheat rhizosphere microbiome: evidences for a core microbiome across eight African and European soils. FEMS Microbiology Ecology 96: fiaa067.

Simonin, M., Préveaux, A., Marais, C., Garin, T., Arnault, G., Sarniguet, A., and Barret, M. (2023) Transmission of synthetic seed bacterial communities to radish seedlings: impact on microbiota assembly and plant phenotype. Peer Community Journal 3:.

Simpson, G.L. (2022) permute: Functions for Generating Restricted Permutations of Data.

Tobin, J. (1958) Estimation of Relationships for Limited Dependent Variables. Econometrica 26: 24–36.

Trivedi, P., Leach, J.E., Tringe, S.G., Sa, T., and Singh, B.K. (2020) Plant–microbiome interactions: from community assembly to plant health. Nat Rev Microbiol 18: 607– 621.

Vandenkoornhuyse, P., Quaiser, A., Duhamel, M., Le Van, A., and Dufresne, A. (2015) The importance of the microbiome of the plant holobiont. New Phytologist 206: 1196– 1206.

Vorholt, J.A., Vogel, C., Carlström, C.I., and Müller, D.B. (2017) Establishing Causality: Opportunities of Synthetic Communities for Plant Microbiome Research. Cell Host & Microbe 22: 142–155.

Weissman, J.L., Hou, S., and Fuhrman, J.A. (2021) Estimating maximal microbial growth rates from cultures, metagenomes, and single cells via codon usage patterns. Proceedings of the National Academy of Sciences 118: e2016810118.

Wickham, H. (2016) ggplot2: Elegant Graphics for Data Analysis, Springer-Verlag New York.

Wippel, K., Tao, K., Niu, Y., Zgadzaj, R., Kiel, N., Guan, R., et al. (2021) Host preference and invasiveness of commensal bacteria in the Lotus and Arabidopsis root microbiota. Nat Microbiol 6: 1150–1162.

Xiong, C., Singh, B.K., He, J.-Z., Han, Y.-L., Li, P.-P., Wan, L.-H., et al. (2021) Plant developmental stage drives the differentiation in ecological role of the maize microbiome. Microbiome 9: 171.

Yan, Y., Kuramae, E.E., Klinkhamer, P.G.L., and van Veen, J.A. (2015) Revisiting the Dilution Procedure Used To Manipulate Microbial Biodiversity in Terrestrial Systems. Appl Environ Microbiol 81: 4246–4252.

Zervas, A., Ellegaard-Jensen, L., Hennessy, R.C., Bak, F., Guan, Y., Horn Herms, C., et al. (2022) Diversity and Structure of Bacterial Communities in Different Rhizocompartments (Rhizoplane, Rhizosphere, and Bulk) at Flag Leaf Emergence in Four Winter Wheat Varieties. Microbiology Resource Announcements 11: e00222–22.

Zhou, X., Wang, J., Liu, F., Liang, J., Zhao, P., Tsui, C.K.M., and Cai, L. (2022) Cross-kingdom synthetic microbiota supports tomato suppression of Fusarium wilt disease. Nat Commun 13: 7890.

