## Supplementary Figures for "Synthetic bacterial community colonizes wheat roots grown in soil and mimics the assembly pattern of a field community"

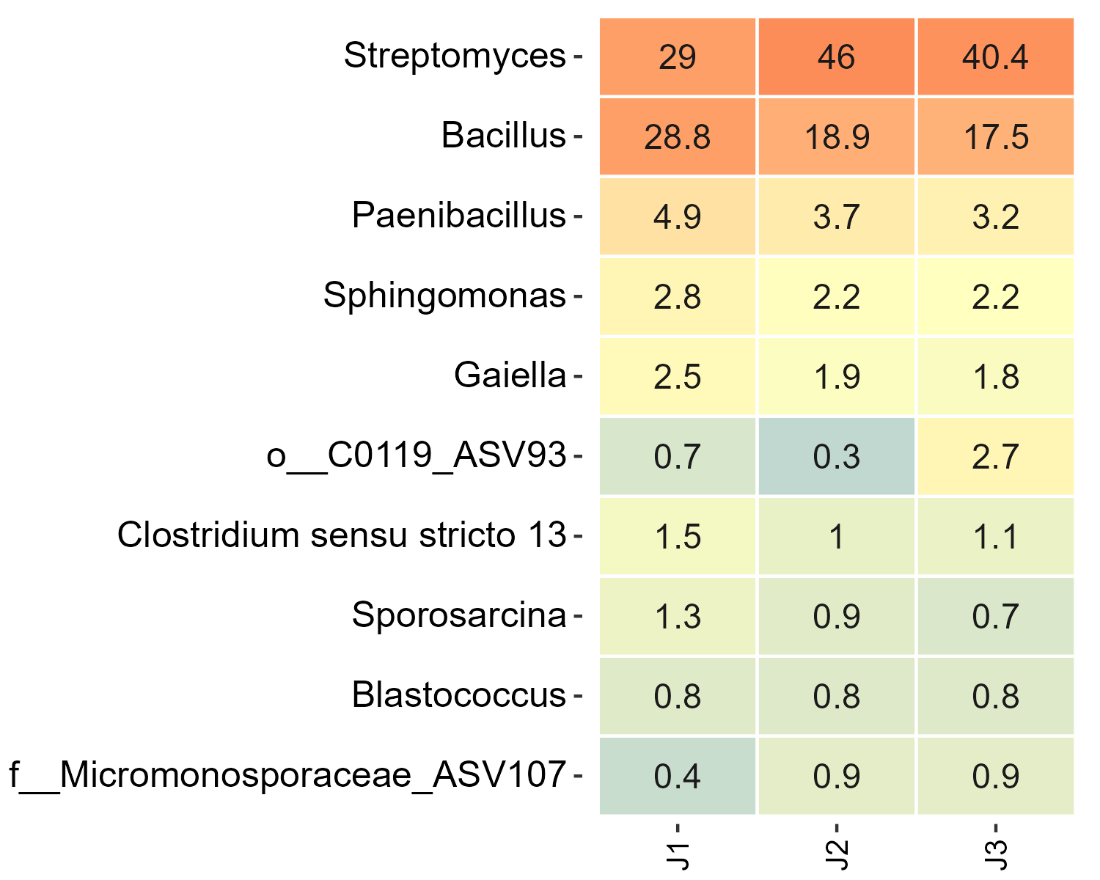

**Supplementary figure 1.** Relative abundance of the 10 most abundant genera in the gamma-irradiated soil.

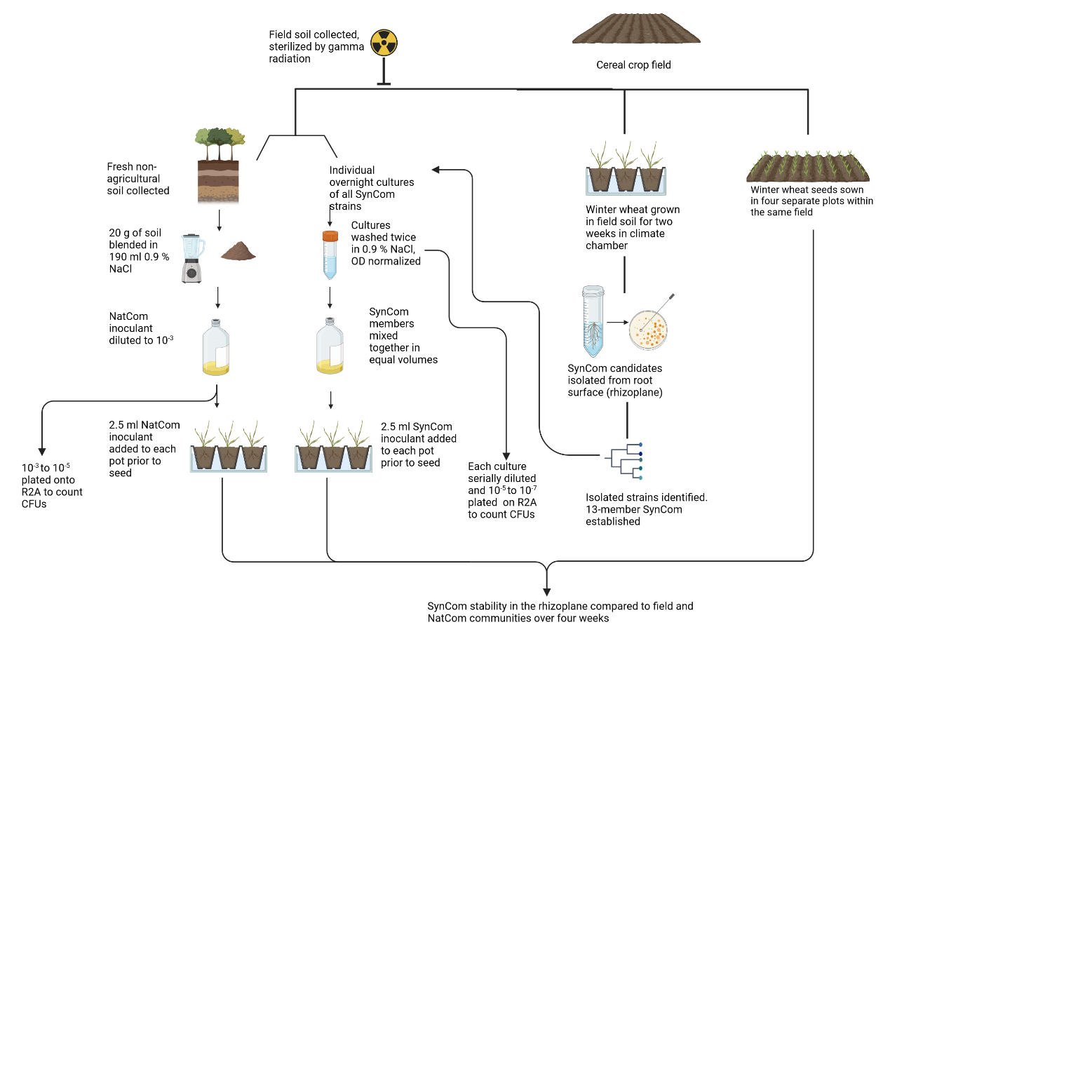

**Supplementary figure S2.** Overview of the experimental design.

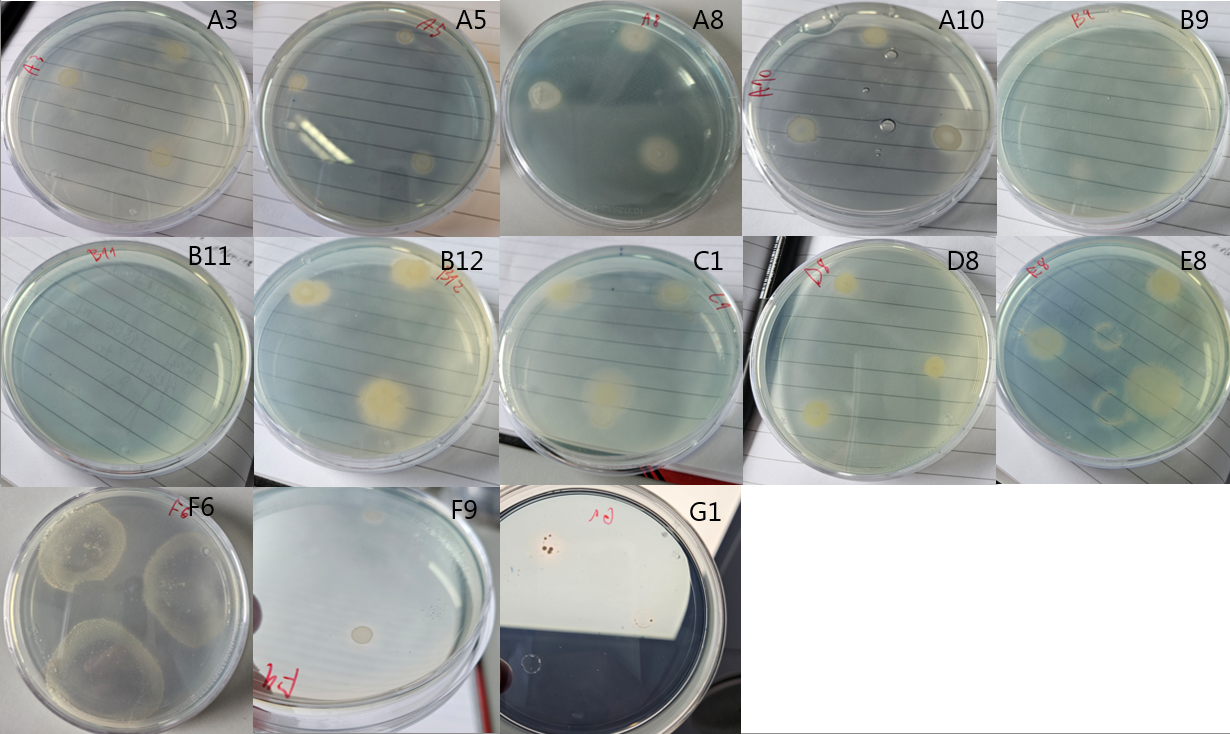
**Supplementary figure S3.** Result of the overlay CAS siderophore assay. Each plate shows one SynCom strain, from top left to bottom right: A3, A5, A8, A10, B9, B11, B12, C1, D8, E8, F6, F9, G1.

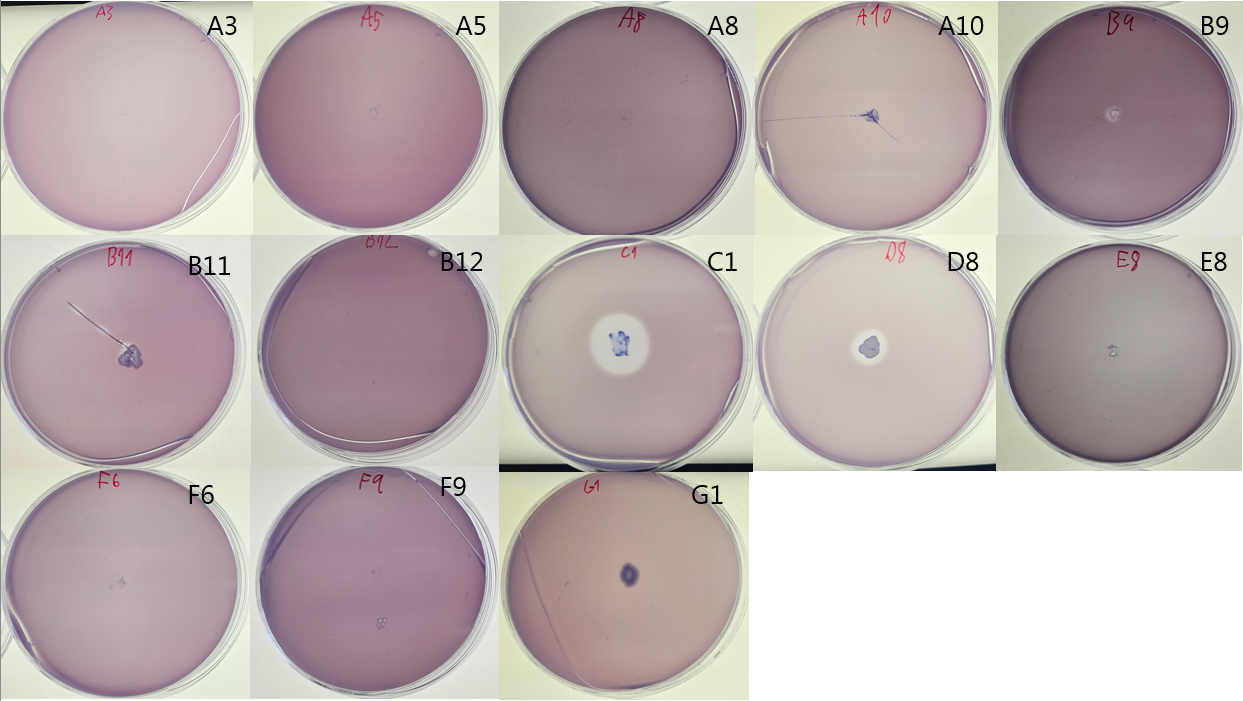
**Supplementary figure S4.** Result of the phytase activity assay. Each plate shows one SynCom strain, from top left to bottom right: A3, A5, A8, A10, B9, B11, B12, C1, D8, E8, F6, F9, G1.

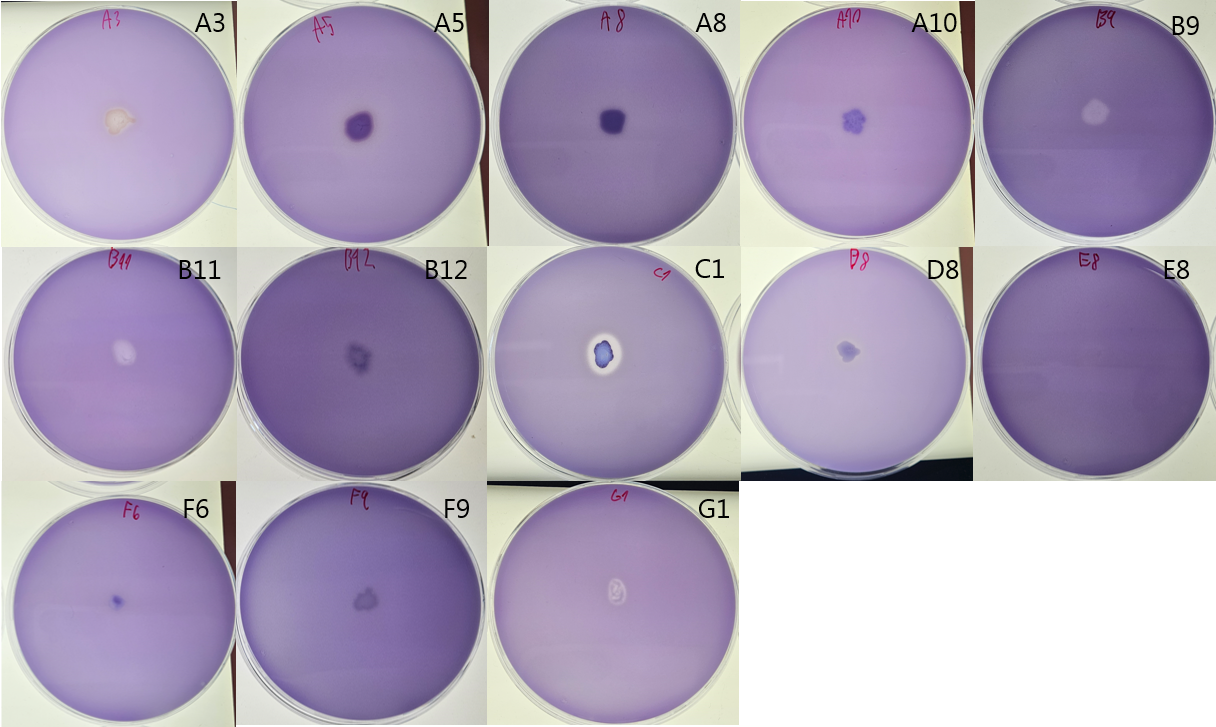
**Supplementary figure S5** Result of the phosphate solubilization assay. Each plate shows one SynCom strain, from top left to bottom right: A3, A5, A8, A10, B9, B11, B12, C1, D8, E8, F6, F9, G1.

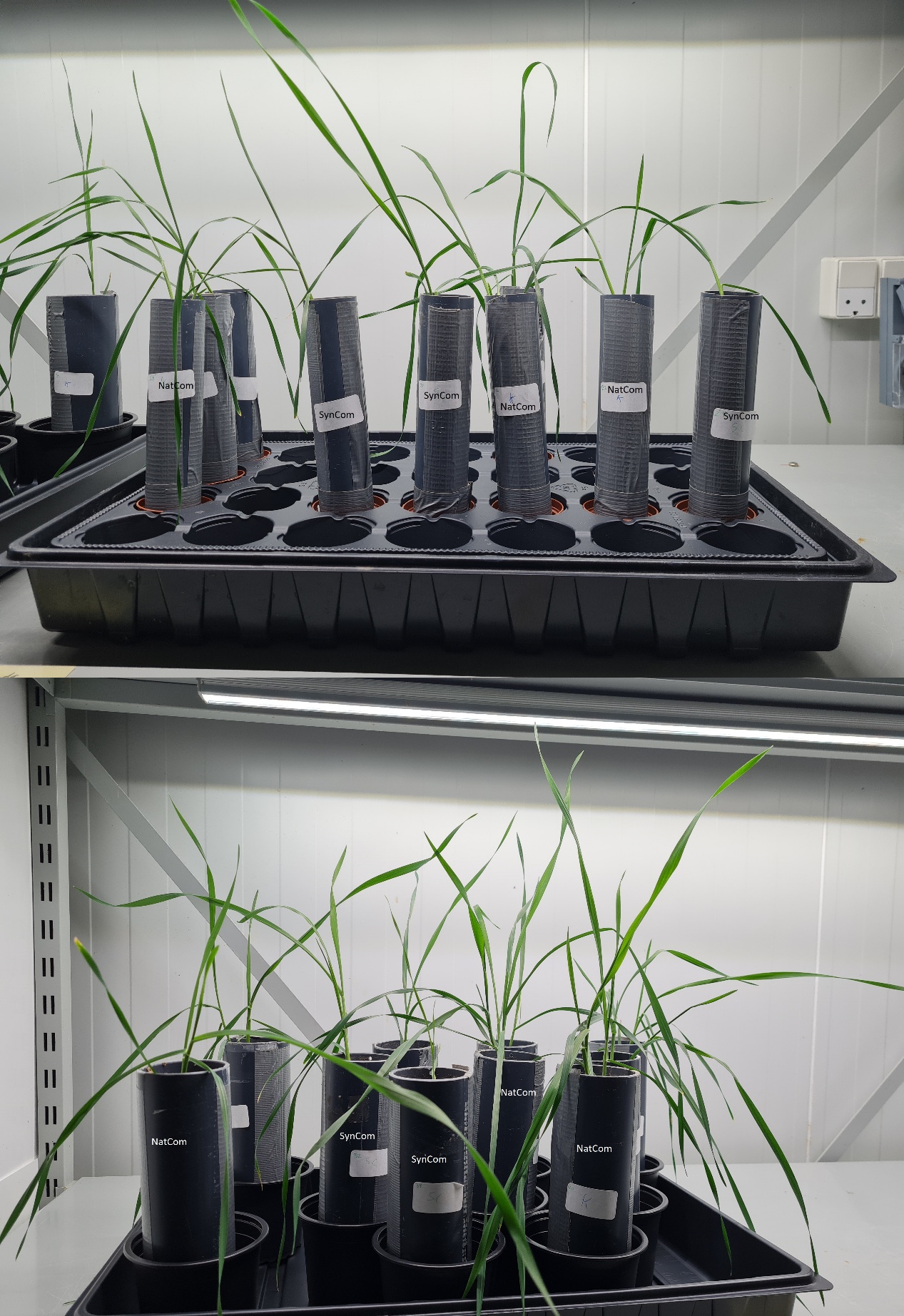

**Supplementary figure 6**. SynCom and NatCom inoculated plants after three (top) and four (bottom) weeks of growth prior to sampling, corresponding to T3 and T4, respectively.

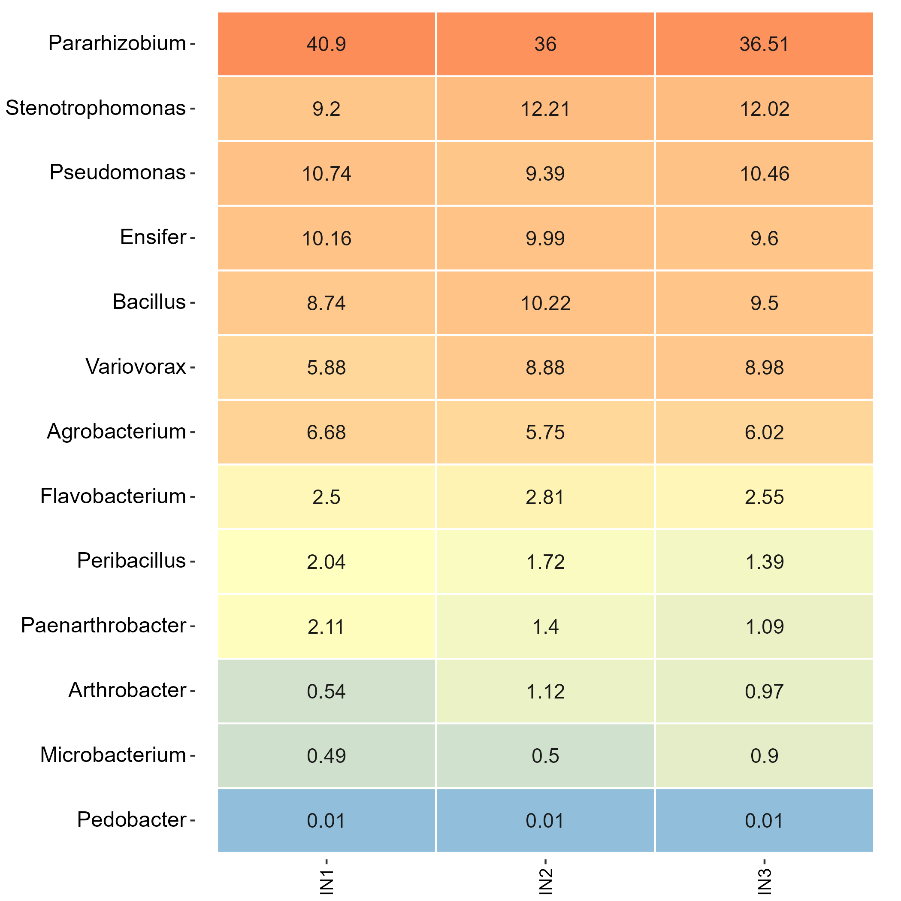

**Supplementary figure 7.** Relative abundances of the 13 SynCom genera in the starting inoculum from three subsamples.

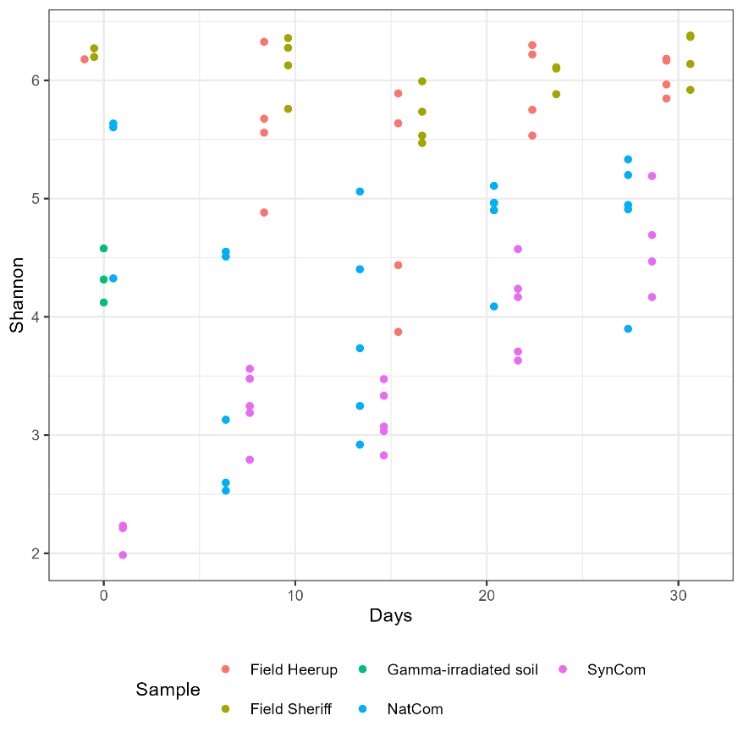

**Supplementary Figure 8.** Shannon diversity. Each point represents a sample. The Shannon diversity was calculated after rarefying the dataset 100 times. At Day 0, the samples from the field are bulk soil samples taken before sowing.

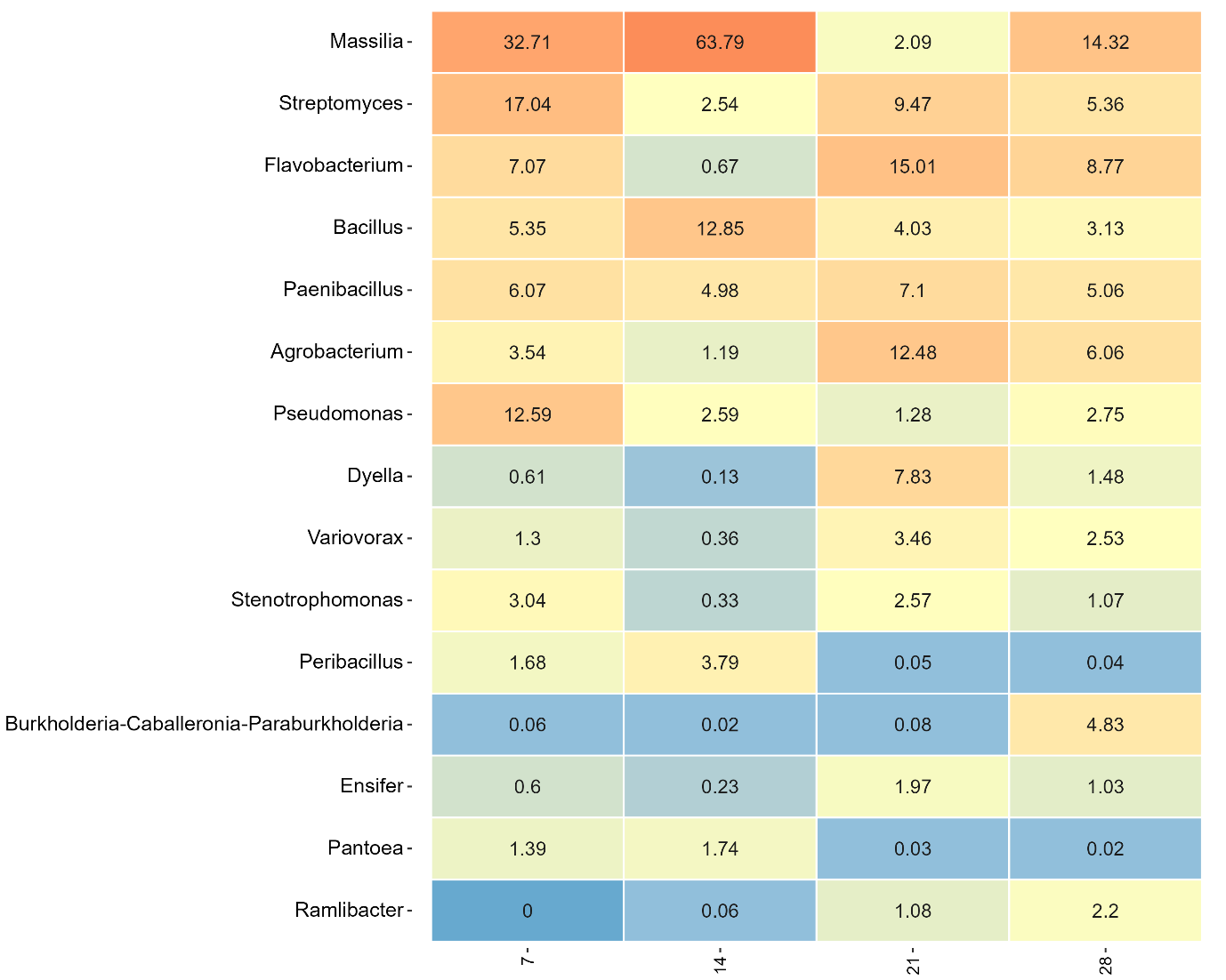

**Supplementary figure 9.** Mean relative abundances of the 15 most abundant genera on the rhizoplane grown in soil inoculated with SynCom strains (n=5). Numbers below the heatmap indicate days after sowing.

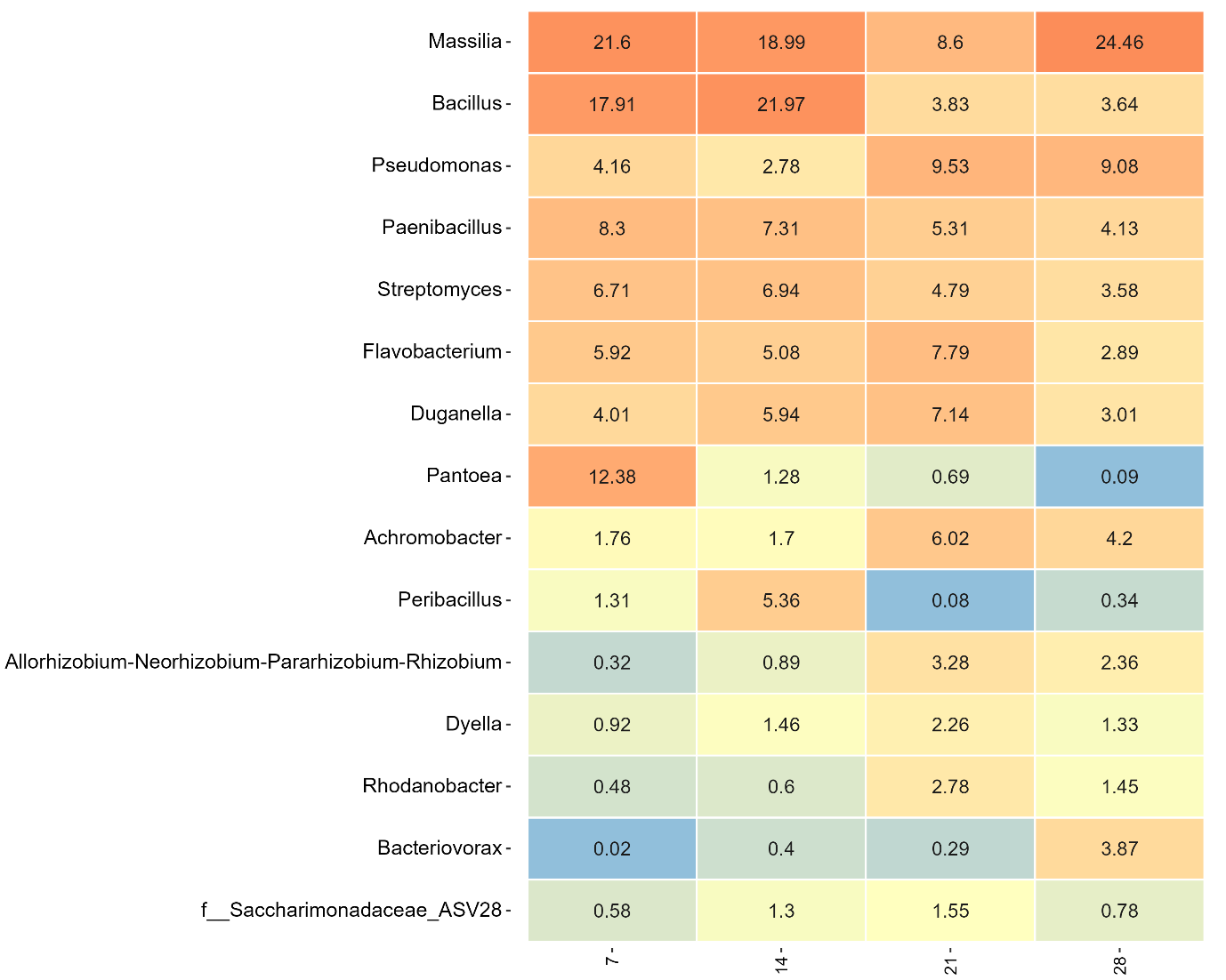

**Supplementary figure 10.** Mean relative abundances of the 15 most abundant genera on the rhizoplane grown in soil inoculated with NatCom strains (n=5). Numbers below the heatmap indicate days after sowing.

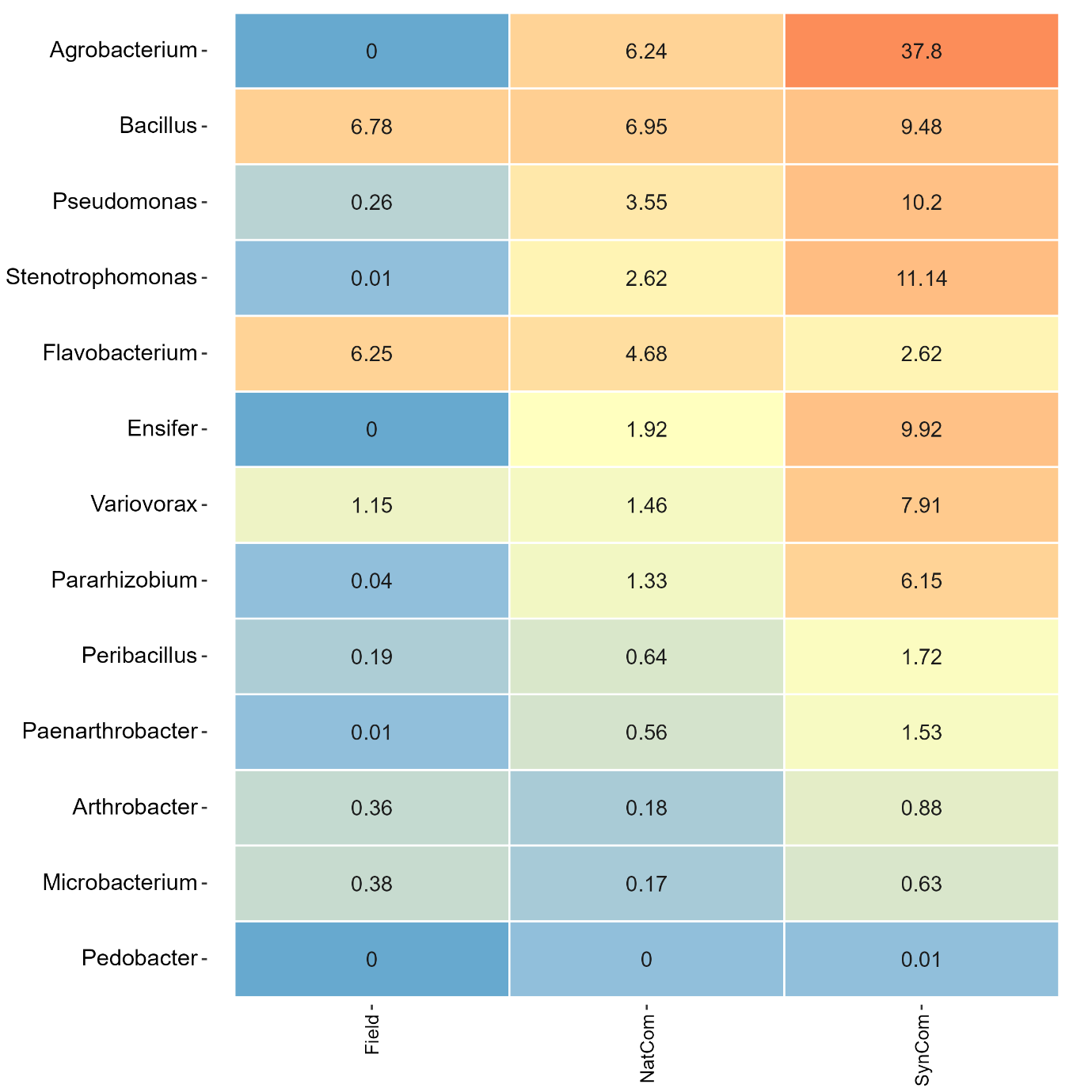

**Supplementary figure 11.** Mean Relative abundances of the 13 genera included in the SynCom, from various starting inocula (n = 3) or bulk soil from the field prior to sowing (n = 3).

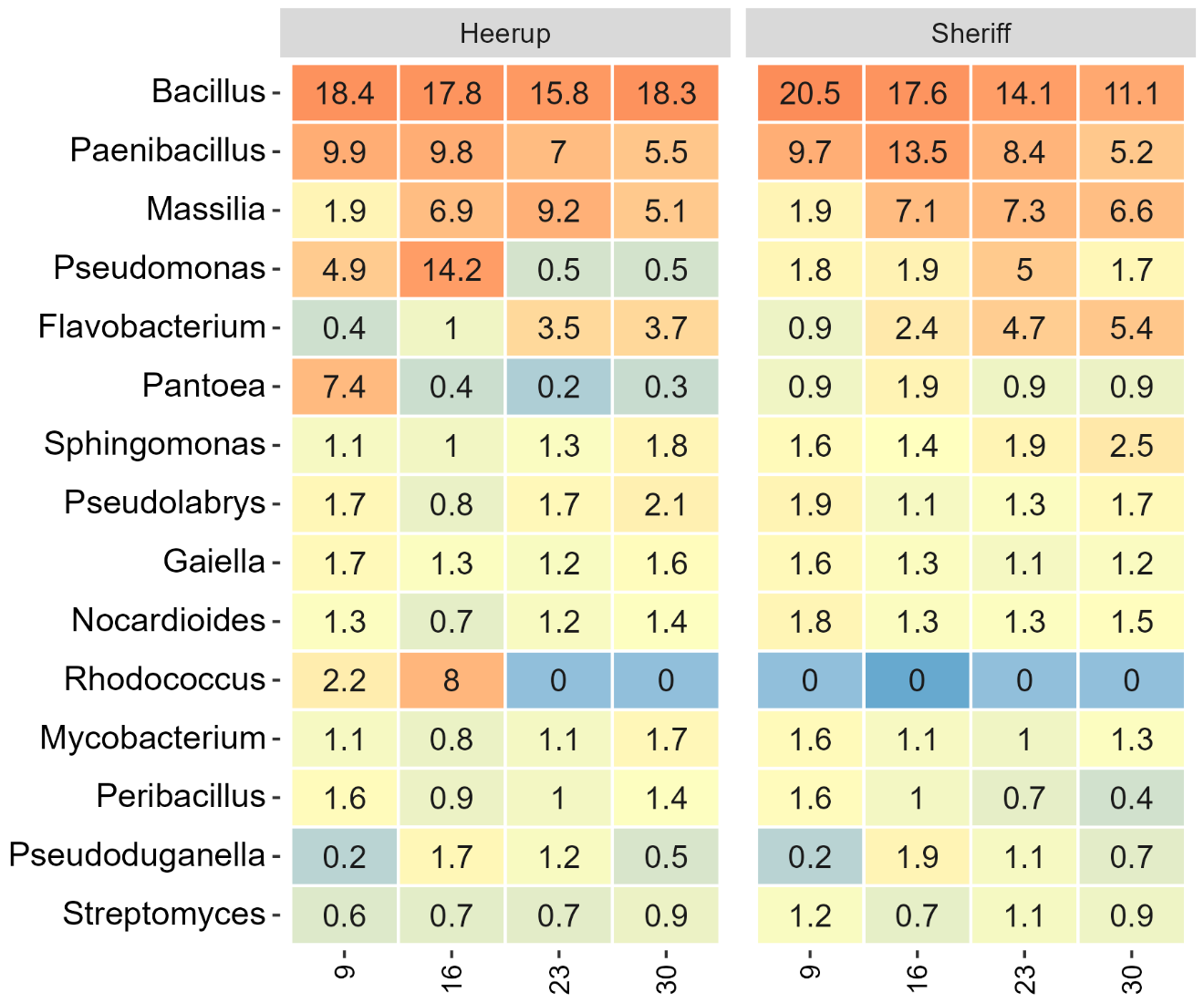

**Supplementary figure 12.** Mean relative abundances of the 15 most abundant genera on the rhizoplane grown in the field of the two winter wheat varieties (n=4). Numbers below the heatmap indicate days after sowing.
