## Supplementary methods for "Synthetic bacterial community colonizes wheat roots grown in soil and mimics the assembly pattern of a field community"

### Isolation of bacterial strains and identification

To isolate rhizoplane bacteria, two grams of each harvested root were rinsed with sterile MilliQ water. Then, the root was suspended in 20 ml of sterile water with 0.85% NaCl and vortexed for 2 minutes. The clean root was subsequently transferred to 20 ml of PBST (0.1 M phosphate buffer, 0.1% Tween 20, pH 7.0) and sonicated for 10 minutes each using an ultrasonic water bath (Struers, Copenhagen, Denmark). This step was repeated twice (transfer and sonication). The roots were removed, and the third suspension was serially diluted, and 100 µl of the 10^-3^ and 10^-5^ dilutions were spread onto 10% R2A (Alpha Biosciences, Baltimore, MD, USA) with 1.5% agar and VL55 agar plates (See below). The plates were incubated for 20 days in the dark at 20°C. Colony color, size, and morphology were used as selection criteria for single colony inoculation into the same medium using 96-well plates containing 150 µl of 10% R2A or VL55 broth, respectively. The 96-well plates were incubated for three days at 20°C and 200 rpm. After incubation, 50 µl of 100% glycerol was added to the 96-well plates to generate cryostocks for a total of 1,344 strains.

To ensure the selection of bacteria that resembles the natural bacterial wheat rhizoplane diversity (Zervas et al., 2022; Guan et al., 2024), we implemented a deselection method using a high-throughput approach. Isolates from each 96-well plate cryostocks were inoculated onto single-well OmniTrays (Thermo Scientific, USA) containing R2A, Gould S1, L9, or R2A supplemented with polymyxin B (10 µg/ml) using a 96-well plate replicator (Sigma-Aldrich, Germany). The Gould’s S1 (Tarnawski *et al.* 2003), L9 (Yim *et al.* 2010), and R2A supplemented with polymyxin B (10 µg/ml) (Van Netten and Kramer 1992) selective media were used to identify the location of *Pseudomonas*, *Sphingomonas*, and *Bacillus* spp. on the 96-well plates, respectively. To identify the rare bacterial genera, we employed an image analysis workflow using the OpenCV library (Version 4.5.3) in Python (Version 3.9). For colony identification, the images of the inoculated OmniTrays were divided into grids corresponding to the 96-well layout. The selected colonies were those that grew on general R2A, but not on Gould’s S1, L9, and R2A with polymyxin B (10 µg/ml). Additionally, we selected 30 strains with unique colony morphology from the Goulds and R2A media supplemented with polymyxin B (10 µg/ml) to isolate *Pseudomonas* and *Bacillus* species. A total of 114 isolates were selected and restreaked on R2A. A single colony from each isolate was inoculated into new 96-well plates containing 150 µl of R2A.

After the deselection process, 114 isolates were selected and re-streaked to ensure strain purity, prior to full length 16S rRNA gene sequencing. First, the gDNA was released from the cells in a 96-well plate format using the lysis method described by (Zhang *et al.* 2021). Subsequently, 2 µl of the resulting bacterial lysate served as a template for PCR amplification of the 16S rRNA gene using the 25 µl 2X Master mix (NZYTech, Lisboa, Portugal) and universal primers 0.2 µM 27F (5'-AGAGTTTGATCCTGGCTCA-3') and 1492R (5'-TACGGTTACCTTGTTACGACTT-3') and water to reach a volume of 50 µl. The thermal cycling conditions used were: initial denaturation step at 97°C for 2 min, followed by 35 cycles of 97°C for 10 s, annealing at 57°C for 30 s, and extension at 72°C for 45 s with a final elongation at 72°C for 10 min. The amplified fragments were cleaned up using the AmPure XP magnetic beads (Beckmann-Coulter, Indianapolis, In, USA) following the manufacturer's protocol. Amplified fragment concentrations were measured using a Qubit (Thermo fisher, Waltham, MA, USA), normalized to ~5ng/µl and then Sanger sequenced (Eurofins, Konstanz, Germany).

### 16S rRNA amplicon data used for strain selection

We downloaded amplicon data from Zervas *et al.* (2022) and Guan *et al.*(2024), which include 16S rRNA amplicon data for the rhizoplane of winter wheat cultivar Sheriff grown for 142 or 14 days, respectively. Two different primer sets were used for the two studies, targeting the V3-V4 or V5-V7 regions in Zervas *et al.* (2022) and Guan *et al.*(2024), respectively. Each data set was processed individually as described previously (Zervas *et al.* 2022; Guan *et al.* 2024).

#### Culture conditions and bacterial strains

An overview of the 13 selected bacterial SynCom strains employed in this study is given in Table S1. The taxonomy of the selected SynCom strains were determined by alignment of their 16S rRNA nucleotide sequences with type strains in the NCBI database using BLASTn.

Strains *Pedobacter* sp. A5 and *Flavobacterium* sp. F6 were maintained on tryptic soya agar (TSA) (Sigma-aldrich, USA), whereas all other strains were maintained on R2A. R2A media contained, per liter: 3 g R2A Broth (Alpha Biosciences, USA) and 15 g bacteriological agar (Millipore, USA).

### Phenotyping of SynCom strains

For all assays, SynCom strains were initially grown in liquid R2A or TSA (Table S1) overnight, then the cells were washed twice in sterile MilliQ water, and the cell suspension adjusted to OD_600_=0.3 to normalize cell densities. Three µl of adjusted cultures were spotted onto the plates, and subsequently incubated at 28°C for 3-21 days depending on the specific assay. Detailed information is provided in Supplementary methods. In brief, the SynCom strains’ abilities to solubilize phosphate was determined by culturing them on National Botanical Research Institute's phosphate growth (NBRIP) medium agar, commonly used to isolate phosphate solubilizing bacteria from environmental samples (Nautiyal 1999). Phytate degradation potential was determined by inoculation on phytate-containing media (Howson and Davis 1983). Clearing zones around colonies on phosphate or phytate media was indicative of phosphate solubilization or phytase activity, respectively. All 13 SynCom members were inoculated onto three plates. We defined a positive result as the formation of a clearing zone >one mm forming around a colony's edge after a maximum of 14 days at all three plates.

### Media for phosphatase and phytase activity

The solid NBRIP(Nautiyal 1999) media contained per liter: 15 g agar, 10 g glucose, 5 g Ca_3_(PO_4_)_2_, 5 g MgCl_2_·6H_2_O, 0.25 g MgSO_4_·7H_2_O, 0.2 g KCl, 0.1g (NH_4_)_2_SO_4_, and 0.005 g bromophenol blue indicator dye. Phytate media was prepared according to Howson & Davies (1983) (Howson and Davis 1983), with some modifications, and contained per liter: 20 g agar, 2.2 g CaCl_2_, 4.96 g NH_4_NO_3_, 0.5 g KCl, 0.5 g MgSO_4_, 0.01 g FeSO_4_, 0.08 g MnSO_4_ and 0.005 g bromophenol blue. After autoclaving, media was cooled to 55°C and10 ml of a 50% w/w phytic acid solution (Aldrich, USA), and 75 ml of a 50% D(+)-glucose solution sterilized with 0.22 µm cellulose acetate filters (VWR, USA) was added to the media. pH was adjusted to 6.0 with sterile 1M NaOH.

### Nitrate reduction

The nitrate reduction assay was performed using the API 20E kit for biochemical identification (BioMerieux Inc, France), and based on the Griess reaction (Tsikas 2007). First, we established cultures of SynCom strains in R2A broth and incubated them overnight. Then, we transferred 200 µl cultures to three ml of nitrate broth (5 g/l peptone (VWR Chemicals, USA), 3 g/l beef extract (Alpha Biosciences, USA) and 1 g/l KNO_3_), and incubated nitrate broth cultures for 24 hours. After incubation, we transferred one ml of nitrate broth culture for each SynCom strain to a sterile 1.5 ml microcentrifuge tube (Eppendorf, Germany) and one drop each of reagents Nit1 and Nit2 (sulfanilic acid and α-naphtylamine, respectively) was added as stated in the protocol. After 5 minutes of incubation at room temperature, formation of a red color indicated a reduction of NO_3_^-^ to NO_2_^-^. The absence of red color is usually indicative of a negative phenotype; however, it can also be caused by reduction of NO_3_^-^ all the way to N_2_ (g). As such, we added a few milligrams of zinc powder to each tube, catalyzing the reduction of any remaining nitrate still present in the tubes. Subsequent formation of a red color confirmed the strain was unable to reduce NO_3_^-^.

### Siderophore production

We determined the capability of the SynCom strains to produce siderophores with the overlay chrome azurol S (O-CAS) method described by Pérez-Miranda *et al.* (2007) with some modifications. After seven days of growth on a dilute growth medium, and subsequent overlay with CAS agar, the formation of a yellow halo on three replicate plates was indicative of siderophore production.

The original CAS-media uses HDTMA. HDTMA is a detergent and therefore toxic to gram-positive bacteria in the SynCom. Therefore we made use of a modified version of the CAS-overlay method by Pérez-Miranda *et al.* (Pérez-Miranda *et al.* 2007). Overnight cultures in R2A broth were washed twice by centrifugation at 8,000 g for five minutes. We resuspended the cultures in sterile MilliQ water and adjusted OD_600_ to 0.3. We added three µl adjusted overnight culture of each SynCom strain onto 1:20 R2A with 100 µM 2.2-bipyridyl agar plates in triplicate and incubated the plates for five days. Twelve ml of overlay CAS agar was then cast on top (per liter: 30.24 g PIPES, 9 g agarose, 72.9 mg HDTA, 60 mg chrome azurol S & 10 ml 1mM FeCl_3_∙6H_2_O), and yellow to orange halo after incubation for one day was indicative of siderophore production. All glassware was washed in 6 *M* HCl to remove trace iron.

### Growth media

**VL55 medium**

VL solidifier

|  | **g l^−1^** | **CAS** |
| --- | --- | --- |
| Phytagel | 32.0 | 71010-52-1 |

Autoclave at 121ºC, 15 min.

VL base medium

|  | **ml** | **CAS** |
| --- | --- | --- |
| 2-(N-Morpholino) ethane sulfonic  acid (MES) | 1.95 g | 4432-32-9 |
| 20mM MgSO_4_.7H_2_O | 10.0 | 10034-99-8 |
| 30mM CaCl_2_.2H_2_O | 10.0 | 10035-04-8 |
| 20mM (NH_4_)_2_HPO_4_ | 10.0 | 7783-28-0 |
| Selenite-tungstate solution | 1.0 |  |
| Trace-element solution SL-10 | 1.0 |  |
| Distilled H_2_O | 460.0 |  |

Autoclave at 121ºC, 20 min.

After cooling the VL base medium to 55ºC, the following items were added: 5 ml of glucose solution, 1 ml of vitamin solution I^c^, 3 ml of vitamin solution II^D^ and 1 ml of cycloheximide (50mg/ml), and 6 ml of 200 mM NaOH. A solution of sterilized 100 mM KOH was used to adjust pH to 5.5. Finally, to avoid bubble formation, the VL solidifying solution was added into the VL base. Further, both solutions should be at the same temperature to avoid clumps forming.

Glucose solution

|  | **g l^−1^** | **CAS** |
| --- | --- | --- |
| Glucose 200mM | 39.67 | 50-99-7 |

Mix well and sterilize by filtering with a pore of 0.22 µm.

Selenite-tungstate solution

|  | **mg l^−1^** | **CAS** |
| --- | --- | --- |
| NaOH | 500.0 | 1310-73-2 |
| Na_2_SeO_3_.5H_2_O | 3.0 | 26970-82-1 |
| Na_2_WO_4_.2H_2_O | 4.0 | 10213-10-2 |

Autoclave at 121ºC for 20 min.

Trace-element solution SL-10

|  | **mg l^−1^** | **CAS** |
| --- | --- | --- |
| HCl (25%) | 10 ml | 7647-1-0 |
| FeCl_2_.4H_2_O | 1500.0 | 13478-10-9 |
| CoCl_2_.6H_2_O | 190.0 | 7791-13-1 |
| MnCl_2_.4H_2_O | 100.0 | 13446-34-9 |
| ZnCl_2_ | 70.0 | 7646-85-7 |
| H_3_BO_3_ | 6.0 | 10043-35-3 |
| Na_2_MoO_4_.2H_2_O | 36.0 | 10102-40-6 |
| NiCl_2_.6H_2_O | 24.0 | 7791-20-0 |
| CuCl_2_.2H_2_O | 2.0 | 10125-13-0 |

Autoclave at 121ºC for 20 min.

Vitamin solution I^c^

|  | **mg l^−1^** | **CAS** |
| --- | --- | --- |
| 4-Aminobenzoate | 40.0 | 555-06-6 |
| (+) - Biotin | 10.0 | 58-85-5 |
| Nicotinic acid | 100.0 | 59-67-6 |
| Hemicalcium  D-(+)-pantothenate | 50.0 | 137-08-6 |
| Pyridoxine-HCl | 150.0 | 58-56-0 |
| Thiamine-HCl | 100.0 | 67-03-8 |
| Cyanocobalamin | 50.0 | 68-19-9 |

Mix well and sterilize by filtering with a pore of 0.22 µm.

Vitamin solution II^D^

|  | **mg l^−1^** | **CAS** |
| --- | --- | --- |
| DL-6,8-Thioctic acid | 10.0 | 1077-28-7 |
| Riboflavin | 10.0 | 83-88-5 |
| Folic acid | 4.0 | 59-30-3 |

Mix well and sterilize by filtering with a pore of 0.22 µm.

#### SynCom plant experiment setup

Wheat plants were grown in PVC pots (height: 19 cm/diameter 3.5 cm or height: 20 cm/diameter 6 cm) as described in Guan *et al*. (2024). PVC pots were rinsed thoroughly in 70% ethanol prior to use. Gamma irradiated soil (sieved 4 mm) was mixed with sand of grain size of 0.4 - 0.9 mm (Dansand, Denmark) in a 3:2 soil:sand ratio. The sand was dried for 24 hours at 105°C and autoclaved twice at 121°C for 15 minutes prior to mixing. We mixed 230 g and 600 g of soil-sand mixture (we refer to this as “soil”) and added it to small and large PVC pots, respectively, for a total of 30 small pots and 10 large pots.

To determine the bacterial communities in the starting soil, we collected one gram of soil from 30 random pots, pooled 10 samples for each replicate (n= 3) in a 50 ml falcon tube and homogenized them by manual shaking for 30 seconds. Afterwards, we transferred 0.5 g from each tube to 1.5 ml Eppendorf tubes and stored them at -20°C until DNA extraction. Soil moisture content was adjusted to 17% w/v with autoclaved MilliQ water. We added 0.79 ml liquid 3-1-4 NPK fertilizer (Park® Drivhusgødning, Germany) to each pot by diluting it in the water used to adjust moisture content. We took three one ml samples from the diluted fertilizer and stored them at -20°C until DNA extraction.

To determine the concentration of live bacteria in the gamma irradiated soil after several years of storage we put aside two grams of soil before any fertilizer addition. We suspended it in 18 ml of sterile 0.9% NaCl for 20 minutes at 300 rpm. A serial dilution from 10^-0^ to 10^-8^ was established with 0.9% sterile NaCl and each concentration spread on 1:2 R2A media in triplicate and incubated at 20°C. After 21 days of incubation, we counted colony forming units (CFUs) on plates averaging 20-200 CFUs.

#### Seed preparation

We surface sterilized seeds of the winter wheat cultivar Sheriff by washing them once in 70% ethanol and once in 5% sodium hypochlorite (Acros Organics, Belgium) for three minutes each wash, followed by three rinses with sterile MilliQ water. Afterwards, we soaked the seeds in sterile MilliQ water for one hour and transferred them aseptically to petri dishes with sterile filter paper, moistened with five ml sterile MilliQ water. The sterilized seeds were left to germinate at room temperature for four days in the dark prior to planting. To verify the efficacy of sterilization 250 µl from the 3^rd^ wash was spread on TSA media and checked for CFUs after one week.

### Isolation of bacterial seed endophytes

To isolate endophytic seed bacteria, two surface sterilized seeds were sown into double autoclaved sand in 50 ml centrifuge tubes aseptically in a LAF bench. The plants were grown at room temperature for 10 days and watered every 3 days using sterile MilliQ water. At day 10, the seedlings were extracted from the sand in a LAF-bench, and the root system was transferred to new 50 ml centrifuge tubes with 20 ml sterile MilliQ water and shaken by hand for 30 seconds to remove excess sand. The roots were then put into 15 ml centrifuge tubes containing 5 mL sterile MilliQ water, and vortexed vigorously for 1 minute. This second wash was serially diluted to 10^-7^, and 100 μl of each dilution was plated onto R2A and TSA, and incubated for up to three weeks at 28°C. Bacterial colonies were picked based on morphology. Isolates were re-streaked until axenic cultures were obtained on R2A, identified through 16S rRNA Sanger sequencing and deposited in glycerol cryostock as described above.

#### Growth chamber experiment

#### SynCom and NatCom inocula preparation

To prepare the SynCom inoculum, we washed overnight cultures of the SynCom strains twice, as described previously for the siderophore inoculation procedure, and adjusted OD_600_ to 0.6 in 0.9% sterile NaCl. We then mixed the washed cells and of the 13 SynCom strains to form the final SynCom inoculum. We put aside 200 µl of each SynCom culture leftover after mixing and serially diluted them up to 10^-7^. Finally, we spread 100 µl of the 10^-5^, 10^-6^ and 10^-7^ dilutions onto TSA plates and incubated them at 28°C and counted the resulting CFUs after 72 hours to verify that live bacteria from all strains had been added and to estimate the cell density for each strain. We collected 500 µl of SynCom inoculum (n=3) and stored at -20°C until DNA extraction for characterization of the starting community.

The NatCom inoculum was created from a native soil microbial community extracted from a non-agricultural soil. First, we collected soil 3-5 cm below the surface two days prior to planting, from the Frederiksberg Campus (55°41'00.5"N 12°32'35.5"E, Frederiksberg, Denmark) and stored it at 5°C until planting. The method used to extract the microbial community followed Yan *et al.* (2015) with a few modifications. Soil was sieved (4 mm) and 20 g was suspended in 190 ml 0.9% NaCl and blended 2 x 2 min. A 1,000-fold dilution was made in 0.9% NaCl to prepare the NatCom inoculum. We took 200 µl of NatCom inoculant for CFU counting as described for the SynCom inoculum. We collected 500 µl of NatCom inoculum (n=3) and stored at -20°C until DNA extraction for characterization of the starting NatCom community.

#### Planting procedure

We dug a two-centimeter-deep hole in the soil in the center of each PVC pot, using sterilized tweezers. Into the holes, we added either 2.5 ml NatCom or SynCom inoculant. Following this, we transplanted one pre-germinated seed to each pot and buried it, before adding one additional milliliter of sterile MilliQ on top of the newly sown seed. The plants were grown for 1-4 weeks in a non-sterile climate chamber with a 16/8-hour day/night cycle at 400 µmol m-^2^ s^-1^ (BX-series LED bars, Valoya, Finland). Temperatures were 20/18.5°C during the day and night, respectively, with a constant 70% relative humidity. We watered the plants every two days as needed, by weighing the entire pots and refilling them with sterile MilliQ up to their original weight at the start of the experiment. The pots were placed in two trays; one for the small pots (30 pots), and one for the large pots (10 pots; see pot sizes above) and every five days the positions of all the pots were randomized within each tray. We sowed 20 plants of each of the SynCom and NatCom treatments, for a total of 40 plants.

### SynCom rhizoplane sampling procedure

We selected five plants grown in small pots from each treatment randomly for sampling of the rhizoplane communities 7, 14, and 21 days after sowing. The plants grown in large pots were sampled 28 days after sowing, with one pot inoculated with SynCom being excluded due to failed germination of the seed. We measured the root and shoot length from the base of the roots. We first washed off the loosely adhering rhizosphere by transferring the roots to 50 ml centrifuge tubes containing 25 ml sterile MilliQ water and shaking by hand for one minute. Subsequently, we collected the soil and bacteria closely adhering to the root surface (rhizoplane) by transferring the roots into a new 50 ml tube with 25 ml sterile MilliQ water, vortexing for one minute and then sonicating for one minute in a water bath. We removed the washed roots from the tubes, dabbed them dry with tissue paper, and measured their wet weight. The tubes with suspended rhizoplane soil were cooled to -80°C for 24 hours and freeze-dried, after which they were stored at -20°C until DNA extraction. All tools and surfaces were cleaned with 70% ethanol between each step of root handling and each individual plant.

### Field sampling

Seven days before sowing, we collected bulk soil from the parcels in each plot with a soil probe rod at a depth of 0 – 20 cm. We sampled five grams of soil from the probe, representative of the entire depth profile sampled. We transferred each sample to a 15 ml tube and homogenized manually by shaking and stored them on dry ice for transport to the lab. Upon arrival, we cooled the bulk soil samples to – 80°C and freeze dried them, before storage at 5°C until DNA extraction.

Nine days after sowing we sampled five plants of both cultivars from each of the four plots at random. We separated the roots and shoots at their base with a sterilized scalpel, and subjected the roots to a two-step washing process. First, we transferred the roots to a 50 ml tube filled with 15 ml sterile MilliQ water and shook them by hand for one minute to remove the rhizosphere soil. We then pooled roots from five plants from each plot, to ensure enough rhizoplane sample for DNA extractions, in a single 50 ml tube with 15 ml MilliQ water and sonicated them for two minutes to dispatch the rhizoplane sample from the roots. We removed the clean roots from the tube with sterile tweezers. We flash-froze the rhizoplane samples immediately in liquid nitrogen and stored them on dry ice. Samples were brought back to the lab and freeze-dried and stored as described for the bulk soil samples. We repeated this sampling procedure at 16, 21 and 28 DAS, except only three plants per treatment were pooled for each sample instead of five, due to larger plants, and the two washing steps were done in tubes with 25 ml MilliQ water. In total, we collected three bulk soil samples prior to sowing and 32 rhizoplane samples (4 time points, 2 cultivars and 4 replicates). All equipment was cleaned with 70% ethanol between each sample. The samples were stored and freeze-dried as described above.

### DNA extraction

Due to the low amount of rhizoplane material for SynCom and NatCom samples at Day 7, 250 µl sterile water was added prior to DNA extraction. For the remaining samples, 100 – 500 mg soil (bulk or rhizoplane) was used for DNA extraction. Samples were homogenized and the cells lysed by bead beating in the FastPrep-24™ 5G system (MP Biomedicals) for 40 seconds at 6.0 m s^-1^. In parallel, three samples containing sigma water were extracted and included as negative controls in the rest of the workflow. After extraction, we stored the DNA at -20°C until further processing.

We assessed the DNA quality with Nanodrop ND-1000 spectrophotometer (Thermo Fisher Scientific, Carlsbad, CA, USA) and further purified it using the NucleoSpin™ gDNA Clean-up mini kit (Machery-Nagel, Germany), according to the manufacturer’s instructions. Following purification, we measured DNA quantity with a QuBit 2.0 Fluorometer (Thermo Fisher) and adjusted samples to 10 ng µl^-1^.

### Library preparation

We prepared the purified DNA for Next Generation Sequencing by performing a first step PCR amplification for library building, using the 779F and 1193R primers (sequences: 5’- AACMGGATTAGATACCCKG -3’ & 5’- ACGTCATCCCCACCTTCC -3’, respectively) (Chelius and Triplett 2001; Bodenhausen, Horton and Bergelson 2013). These primers amplify the V5-V7 regions of the bacterial 16S rRNA gene, with a lowered amplification of contaminating plastid sequences from the host plant compared to other, similar primers (Beckers *et al.* 2016). The 5’ end of the 799F and 1193R primers additionally contained the universal TruSeq adaptors required for sequencing on the Illumina Sequencing platform: 5’- ACACTCTTTCCCTACACGACGCTCTTCCGATCT 3’ and 5’- GACTGGAGTTCAGACGTGTGCTCTTCCGATCT 3’, respectively. We used ZymoBIOMICS Microbial Community DNA Standards (Zymo Research, USA) at this step of the process at identical concentrations. The mock communities contained known ratios of DNA from eight species of bacteria and two fungi with a G-C content varying from 15 to 85% and was used to assess biases or errors introduced during sequencing. Negative controls were included during DNA extraction and library preparation.

We performed PCR amplification in 20 µl reactions composed of 10 µl Platinum II Hot-Start PCR Master Mix (2X) (Thermo Fisher Scientific, USA), 0.5 µl 10 m*M* of 799F & 1193R primers, 0.5 µl 0.3 m*M^1^* bovine serum albumin (New England Biolabs, United Kingdom), 0.5 µl 25 m*M* MgCl_2,_ (Sigma-Aldrich, USA) 5 µl sigma water and 3 µl sample DNA. PCR cycling had the following conditions: Initial 95°C annealing for 2 minutes, 33 cycles of denaturation at 95°C for 15 s, annealing at 55°C for 15 s and 72°C extension for 15 s, and finally 72°C for 5 minutes for the final elongation. The PCR products were kept at 5°C until further use. We verified the quality of the PCR products with QuBit and gel electrophoresis on 1 % agarose gels at 100 mV for 20 minutes. In the final step before sequencing, we did an additional purification with the AMPure XP beads (Beckman Coulter Life Sciences, USA), added in a ratio of 12:15 of PCR-product:bead volume. We placed the mixture onto a magnetic rack, washed twice with 100 µl 80% ethanol and eluted in 27 µl sigma water to obtain pure PCR products.

Amplicon processing was performed in dada2 as in Guan *et al.* (2024), with a few modifications. Primers were removed using trimLeft. In the filtering step, maxEE was set to (2,2), and forward and reverse reads were truncated to 280 and 200 bp based on quality inspection, respectively.

### References

Beckers B, Op De Beeck M, Thijs S *et al.* Performance of 16s rDNA Primer Pairs in the Study of Rhizosphere and Endosphere Bacterial Microbiomes in Metabarcoding Studies. *Frontiers in Microbiology* 2016;**7**, DOI: 10.3389/fmicb.2016.00650.

Bodenhausen N, Horton MW, Bergelson J. Bacterial Communities Associated with the Leaves and the Roots of Arabidopsis thaliana. *PLOS ONE* 2013;**8**:e56329.

Chelius MK, Triplett EW. The Diversity of Archaea and Bacteria in Association with the Roots of Zea mays L. *Microbial Ecology* 2001;**41**:252–63.

Guan Y, Bak F, Hennessy RC *et al.* The potential of Pseudomonas fluorescens SBW25 to produce viscosin enhances wheat root colonization and shapes root-associated microbial communities in a plant genotype-dependent manner in soil systems. *mSphere* 2024;**0**:e00294-24.

Howson SJ, Davis RP. Production of phytate-hydrolysing enzyme by some fungi. *Enzyme and Microbial Technology* 1983;**5**:377–82.

Nautiyal CS. An efficient microbiological growth medium for screening phosphate solubilizing microorganisms. *FEMS Microbiology Letters* 1999;**170**:265–70.

Pérez-Miranda S, Cabirol N, George-Téllez R *et al.* O-CAS, a fast and universal method for siderophore detection. *Journal of Microbiological Methods* 2007;**70**:127–31.

Tarnawski S, Hamelin J, Locatelli L *et al.* Examination of Gould’s modified S1 (mS1) selective medium and Angle’s non-selective medium for describing the diversity of Pseudomonas spp. in soil and root environments. *FEMS Microbiology Ecology* 2003;**45**:97–104.

Tsikas D. Analysis of nitrite and nitrate in biological fluids by assays based on the Griess reaction: appraisal of the Griess reaction in the L-arginine/nitric oxide area of research. *J Chromatogr B Analyt Technol Biomed Life Sci* 2007;**851**:51–70.

Van Netten P, Kramer JM. Media for the detection and enumeration of Bacillus cereus in foods: a review. *International Journal of Food Microbiology* 1992;**17**:85–99.

Yan Y, Kuramae EE, Klinkhamer PGL *et al.* Revisiting the Dilution Procedure Used To Manipulate Microbial Biodiversity in Terrestrial Systems. *Appl Environ Microbiol* 2015;**81**:4246–52.

Yim M-S, Yau YCW, Matlow A *et al.* A novel selective growth medium-PCR assay to isolate and detect *Sphingomonas* in environmental samples. *Journal of Microbiological Methods* 2010;**82**:19–27.

Zervas A, Ellegaard-Jensen L, Hennessy RC *et al.* Diversity and Structure of Bacterial Communities in Different Rhizocompartments (Rhizoplane, Rhizosphere, and Bulk) at Flag Leaf Emergence in Four Winter Wheat Varieties. *Microbiology Resource Announcements* 2022;**11**:e00222-22.

Zhang J, Liu Y-X, Guo X *et al.* High-throughput cultivation and identification of bacteria from the plant root microbiota. *Nat Protoc* 2021;**16**:988–1012.
